# Untying Rates of Gene Gain and Loss Leads to a New Phylogenetic Approach

**DOI:** 10.1101/2025.01.27.634999

**Authors:** Yoav Dvir, Sagi Snir

## Abstract

The advent of the genomic era has produced an incredible wealth and resolution of molecular data, posing an unprecedented challenge for molecular systematics, necessitating novel techniques and paradigms. Consequently, whole genome approaches were developed to extract the evolutionary signal by taking advantage of a larger amount of data. In parallel and in light of the understanding that in prokaryotes, genome dynamics (GD) events, primarily gene gain and loss, provide a significantly richer signal than point mutations in ubiquitous housekeeping genes, GD-based approaches were suggested. However, proper modeling of these data and the processes generating them has lagged in their pace of accumulation, both because of a lack of deep understanding and because of technical difficulties. Among the central hurdles of accurate modeling of real data is the relaxation of rate constancy, particularly the untying of gain and loss rates. This relaxation violates key assumptions such as constant genome sizes, gene set, and model reversibility and has vast implications for implementation. This work presents a generic stochastic model, the two-ratio process (TRP), which encompasses and deals with these complications. As a special case, it contains the Poissonian process with different gene gain and loss rates as a form of the Birth-Death process with varying population sizes. The lack of reversibility invalidates traditional phylogenetic approaches, yielding a novel two-stage phylogenetic approach in which accurate, bidirectional parameters are first inferred for triplets and later combined by a special cherry-picking method to a complete tree. We show by algebraic techniques that this method is theoretically statistically consistent. The method implemented by the software TDDR (Triplets Directed Distances Reconstruction) was applied to synthetic data, showing an advantage over other approaches handling similar data but without the same model assumption. We also applied it to the Alignable Tight Genomic Clusters (ATGC) Database, which showed a high adequacy to the observed data. The TDDR code is available on GitHub: https://github.com/YoavDvir/TDDR.

## 1 Introduction

Modeling genomic processes presents significant challenges on multiple fronts [24,15]. From a statistical perspective, we must accurately characterize key factors and their contributions while maintaining the simplicity of the model [4,3,18]. Computationally, most likelihood-based approaches are intractable [12,52], necessitating simplified yet accurate system abstractions or sound heuristics [65,25,30]. From a biological point of view, as noted in [79], “Biological systems reach hierarchical complexity that has no counterpart outside the realm of biology.”

These challenges are particularly evident in phylogenetics - the task of reconstructing the evolutionary history of organisms. Given the unprecedented availability of high-quality genomic sequences, addressing these challenges has become crucial [31]. Phylogenetics represents one of the more complex biological processes [63], with horizontal (or lateral) gene transfer (HGT) being a principal obstacle to accurate analysis, especially in prokaryotes. HGT, the transfer of genetic material through non-vertical inheritance routes [16,36,47], is prevalent in prokaryotes, connecting distant lineages in the Tree of Life and effectively transforming it into the Network of Life [42,51,77]. However, since most genetic material follows parent-child inheritance, the signal of vertical tree-like evolution remains discernible even among prokaryotes [1,50,83].

This situation requires phylogenetically sensitive markers capable of distinguishing even between species strains. Traditional phylogenetic markers, primarily ribosomal or other housekeeping genes [19], while robust to HGT and ubiquitous, often lack sufficient sensitivity [38]. Within prokaryotes, genome dynamics (GD), the relentless mobility of DNA between organisms, has emerged as a promising solution [64,56,48,37]. GD encompasses three primary operations: duplication, transfer, and loss (DTL) [7,66,46,72,17], with transfer and loss accounting for approximately 98% [49]. GD-based phylogenetics divides into order-based approaches [55,27,80], using gene order along chromosomes as markers, and content-based methods [64,22], which analyze the symmetric difference between gene sets. The synteny index [62,2] combines both approaches by comparing neighborhoods of common genes.

GD typically creates significant discordance between individual gene trees showing single-gene evolution and the species tree representing the main evolutionary trend at the entire species level. This has spawned two related research directions: reconstructing species trees from gene trees (*supertree* construction) [14,68,43,53,26,28], and *gene-tree/species-tree reconciliation* by accounting for DTL events of this gene along the given species tree [41,82,6,5]. Both approaches construct gene trees using orthologous sequences (i.e., COGS [69]), either DNA or protein, from various species.

The increasing prominence of machine learning in genomics has made accurate modeling essential. Time-dependent models are needed to define events and their probabilities, enabling likelihood-based approaches that link observations with likely event sequences [21,5,53,28]. Several GD-based phylogenetic models exist, including genome rearrangement [59,55,9,67] and gene gain/loss [30,78,71,77,23]. A major challenge lies in distinguishing between gain and loss rates, which challenges the fundamental phylogenetic concept of *model reversibility* – model independence from time direction [70]. Notable work by [30] presents a simple birth-and-death process with apparently independent rates, though the assumption of equilibrium enforces a constant genome size and thus reversibility. An earlier study [32] proposed a birth-death-innovation process with independent genome rates, although its framework lacks direct phylogenetic application. [77] describes a symmetric gene content-based approach (SGC) that, while lacking a well-defined model, serves as an evolutionary “clock” for prokaryotic GD. The synteny index analysis relies on a single operation model, the Jump [61,60,34].

Our work takes a crucial step forward in GD-based phylogenetics by establishing theoretical foundations to separate gain and loss rates in bacterial evolution, a property supported by microbial studies [57]. We propose the two-ratio process (TRP), a framework independent of gene order, which encompasses various models, including order-based, Poissonian birth-and-death and symmetric unidirectional distance-based approaches. Focusing on prokaryotes, our model excludes duplication-based gains due to their negligible frequency compared to HGT (see, e.g., [75,73,36,49]). Another key assumption of the model underlying the TRP is the model of the *infinite pangenome* (or *supergenome*) model, accounting for the vast prokaryotic gene repertoire [13,49] and gene set diversity even within a single species [76].

We demonstrate the model’s additivity [58], allowing path-based measurements, and present an efficient polynomial-time algorithm, the triplet directed distance-based reconstruction Algorithm. TDDR operates on gene sets on leaves, transforming TRP modeling into bidirectional routes in the tree. Using algebraic arguments, we show that triplets, rather than pairs of species, are necessary despite the increase in computational complexity, and we prove the statistical consistency of TDDR [21].

We implemented TDDR and evaluated it against existing software using simulations and real-life data sizes. TDDR demonstrated superior performance in topology reconstruction and parameter estimation. Analysis of bacterial families from the Alignable Tight Genomic Clusters (ATGCs) database [39] revealed a significant variance between gain and loss rates, confirming the importance of rate disentanglement. The TDDR software, implemented in Python, is publicly available at https://github.com/YoavDvir/TDDR. The appendix includes proofs, a broader description of the theory, and experimental results. To enable fluent reading, the sections of the Appendix are extended versions of correspondent sections of the main body.

## 2 The Model

(For an extended version of this section, see Sect. A)

In this part, we introduce the framework that accounts for different and changing gain and loss rates and underlies this entire work. We describe a stochastic process in which a root genome evolves through the binary-rooted phylogenetic tree to leaf genomes. Our building block is the two-ratio process (TRP), which represents a stochastic process that operates along the edges and paths of the tree *T*. Throughout this work, we assume that a genome is a set of unique genes (i.e., a single copy of every gene). A central underlying property used throughout the work is the *infinite pangenome model* [13,49], which implies that a given gene can be gained only once throughout history, maintaining the uniqueness of the gene in a genome. The following lemma, which we denote by *the convexity lemma* [45,44], follows directly from the infinite pangenome.

### Lemma 1.

*The set of genomes containing a specific gene forms a connected component in the underlying tree T*.

It is also assumed that the TRP operates independently and individually along the directed edges. A TRP denoted by 𝒢_0,1_, has tw o parameters 0 *< R*_0,1_, *R*_1,0_ *<* 0 (the ratios). 𝒢_0,1_ starts with a parent genome 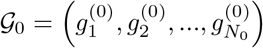, with *N*_0_ different genes, which evolves to a child genome 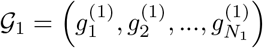 with *N*_1_ different genes. A gene in 𝒢_0_ *survives* to be in 𝒢_1_ with probability *R* 0,1. Each gene 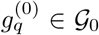 is replaced in 𝒢_1_ by a set *M*_*q*_ of genes, possibly including 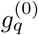 itself. All genes in *M*_*q*_ except 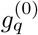 are new. Denote by *X*_*q*_ the number of genes in *M*_*q*_. It is assumed that 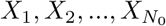 are i.i.d. with expectation *e*_0,1_ and variance *v*_0,1_. We define 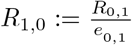. The variance is later considered to be a nuisance parameter. Fig. 1a illustrates the TRP. The number of genes that appear in both 𝒢_0_ and 𝒢_1_ is denoted by *N*_0,1_.

**Fig. 1.**
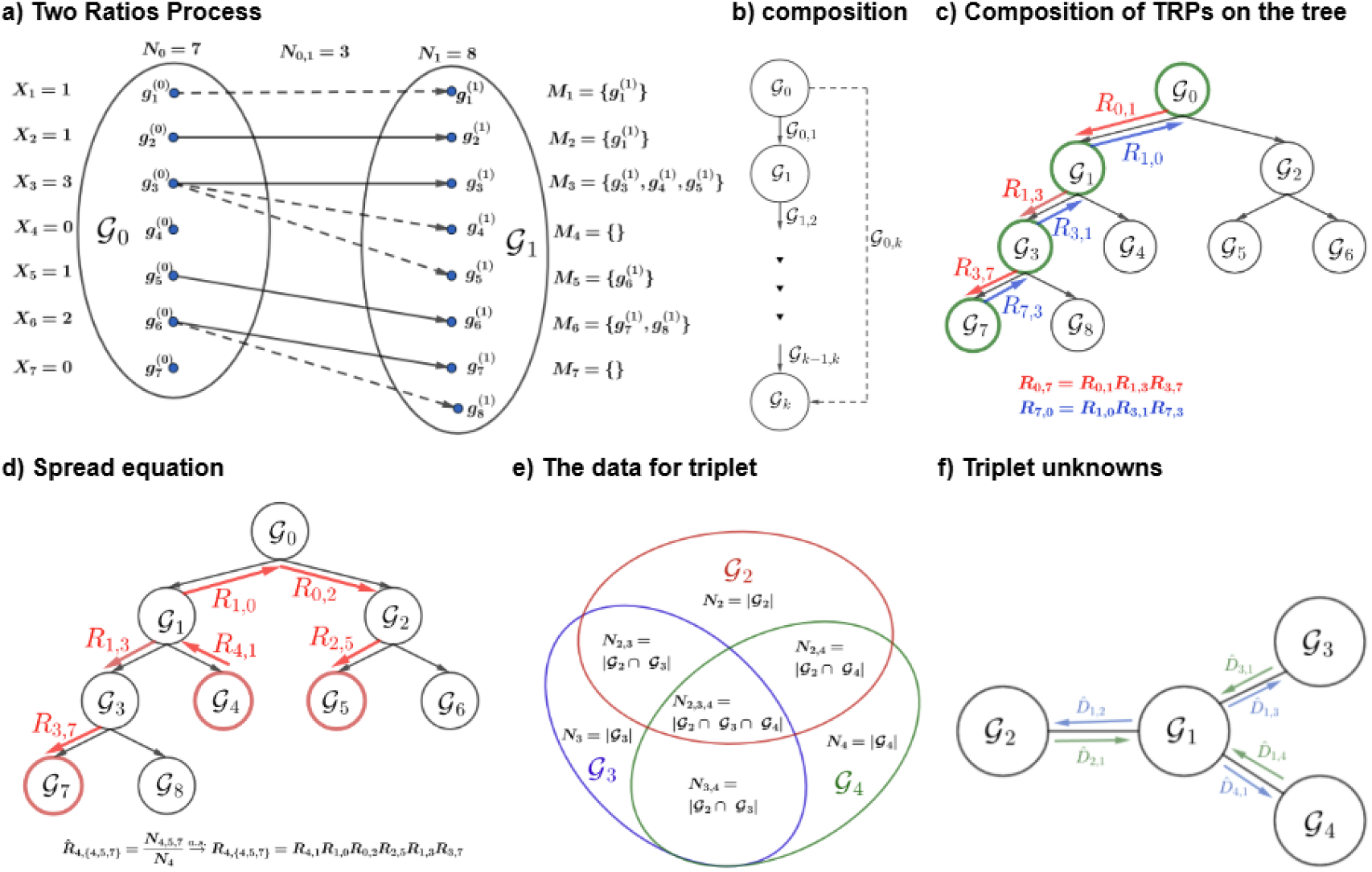
a) Two Ratio Process: The genome 𝒢_0_ evolves to genomes 𝒢_1_. The size of 𝒢_0_ is *N*_0_ = 7. The size of 𝒢_1_ is *N*_1_ = 8. Every gene 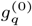 in 𝒢_0_ is replaced by a sequence *M*_*q*_ with a length greater or equal to zero. The gene 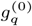 may appear in 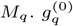 cannot appear *M*_*j*_, *j ?*≠ *q*. An arrow goes from 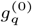 to the genes of the set *M*_*q*_ in *G*_1_. An arrow with the continuous line goes from a gene in 𝒢_0_ to the same gene in 𝒢_1_. The number of genes that appear both in 𝒢_0_ and in 𝒢_1_ is *N*_0,1_ = 3. **b) Composed Process of several TRPs**: The genome *G*_*i*_ evolves to genomes 𝒢_*i*+1_ by the TRP 𝒢_*i,i*+1_. The composed process from 𝒢_0,*k*_ from 𝒢_0_ to 𝒢_*k*_ is also a TRP. **c) Composition of TRPs on a Phylogenetic Tree**: There is TRP for each edge of the tree from parent to child. For example 𝒢_1,3_ from 𝒢_1_ to 𝒢_3_. Due to the composition the process 𝒢_0,7_ is also a TRP with ratios *R*_0,7_ and *R*_7,0_. 𝒢_1,7_ and 𝒢_0,3_ are also TRPs. Note the direction of the TRP is always down the tree. 𝒢_7,0_ is not a TRP. **d) Spread equation**: The spread ratio *R*_4,*{*4,5,7*}*_ from 𝒢_4_ to 𝒢_5_ and 𝒢_7_ is defined by the multiplication of the edge ratios that are notated by the red arrows. An estimator for this spread ratio is 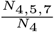.Applying the −log function, we obtain 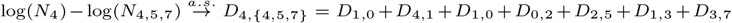. Thus, the spread equation is 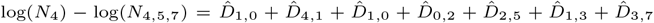.**e) The data of three leaves**: the data of three leaves consist of the genome sizes, the sizes of the intersections of pairs of leaf genomes, and the size of the intersection for all triplets of leaf genomes. **f) The edge-directed distances are the unknowns:** The edge-directed distances from 𝒢_1_ to the leaves and the directed distances from the leaves to 𝒢_1_ are the unknown parameters that need to be estimated. There is a linear transformation with a kernel of dimension one between the vector of logs of the sizes on the left and the vector of the unknowns that appear on the right. If the size of the intersection of the three genomes is omitted, then there is not enough data to obtain the estimates.

### Observation 1 (TRP Properties:)

*If N*_0_ *tends to infinity, then by the strong law of large numbers, we have*

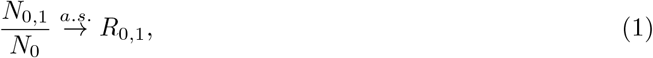

*Applying the strong law of large numbers, we again deduce that* 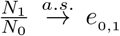. *Combining that with the previous item*,

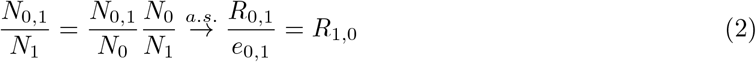

*In words, it means that* 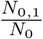 *and* 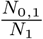 *tend almost surely to R*_0,1_ *and R*_1,0_ *respectively*.

From these properties, it follows that 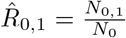 and 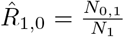 are estimators for the parameters *R*_0,1_ and *R*_1,0_.

**Comments:**

a. Although *R*_0,1_ is a probability, the ratio *R*_1,0_ is not a probability, and there is no reversed TRP process 𝒢_1,0_.
b. Later, when we deal with unrooted phylogenetic trees, the edges of the tree are TRPs, and it is not known which of the two genomes starts and which ends the process. Since, at least in one direction, the ratio is not a probability, we prefer to refer to both as ratios.

### 2.1 Birth-and-Death Process as a Special Case of the TRP

As mentioned above, the TRP model has, as a special case, several more popular models. Here, we describe one.

#### Observation 2

*If it is assumed that* 𝒢_0_ *evolved to* 𝒢_1_ *by a linear birth-and-death process [20, p. 456] n*_*t*_, 0 ≤ *t* ≤ *τ where τ* = *−*(*logR*_0,1_ + *logR*_1,0_), *with a birth rate* 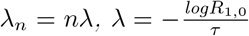 *and a death rate* 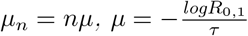., *then it is possible to describe such a process between genomes as a TRP*.

The appendix gives a formal and detailed description in subsection A.1.

### 2.2 Composition of Two or More TRPs

A process composed of two or more TRPs is a TRP (see Fig. 1b). This property is essential for this work and is one of the advantages of the TRP model. It relies on the Convexity Lemma 1 defined above, guaranteeing that the genomes harboring a given gene form a connected subtree. For example, a composition of two or more birth-and-death processes with possibly different parameters is not necessarily a birth-and-death process. One can compose two nonhomogeneous birth-and-death processes, and the result is again a nonhomogeneous birth-and-death process. However, the description of the TRP is more general and straightforward. Section A.2 gives a formal and detailed description in the appendix.

### 2.3 The Phylogenetic Tree

Let *T* = (*V, E*), where *V* = *{*𝒢_0_, 𝒢_1_, …, 𝒢_2*m−*2_*}*, be a connected rooted binary phylogenetic tree with *m* leaves. The convention here is that 𝒢_0_ is the root of the tree. The leaves represent known genomes because their sequence of genes is known, while the internal nodes represent unknown genomes. The input to the model is the size of the gene set in every leaf genome and the intersection sizes among the members of pairs or triplets of the leaves.

Let *S* be a set of genome indexes. Denote the number of genes all these genomes share by *N*_*S*_. We may omit parentheses of the set in the notation. Thus, we denote *N*_2,4_ instead of *N*_*{*2,4*}*_.

Then, if *i*_1_, *i*_2_, …, *i*_*m*_ are the indexes of the leaves, the data we use for our estimations are the genome sizes of the leaves, 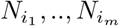, the sizes of pairwise intersections of the leaves, 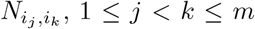, and the sizes of intersection of triplets of genomes, 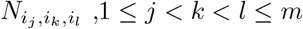.

For each directed edge, with neighboring node genomes 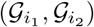, for which 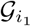 is considered a parent and 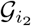 is its child, it is assumed that 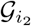 evolved from its parent through a process denoted by 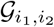. This process acts according to the TRP model described in the previous section. Due to the composition property, even if *i*_2_ is just a descendant of *i*_1_ and not a direct child, then 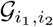 is also a TRP (see Fig. 1c). In the following subsections, we describe a method for inferring the tree structure given leaf genomes assumed to have evolved via TRPs.

### 2.4 Spread Ratio

Given a set *S*_0_ of genome indexes at the nodes of *T*, let 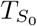 be the minimal connected subtree of *T* that contains *S*_0_, and let 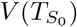 and 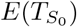 be the set of nodes and edges in 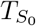, respectively. Now, for *i*_0_ ∈ *S*_0_, let *T*′(*S*_0_, *i*_0_) be the tree 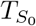 that is rooted in *i*_0_ ∈ *S*_0_, such that the (directed) edges (*a, b*) in *E*(*T*′(*S*_0_, *i*_*o*_)) are directed away from *i*_0_, that is, node *a* is closer to root *i*_0_ than *b*.

The above structure of *T*′(*S*_0_, *i*_0_) allows us to define a new notion that will be essential later.

#### Definition 1.

*For a set S*_0_ *of nodes in T and a node i*_0_ ∈ *S*_0_, *let the spread ratio* 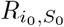 *from i*_0_ *to S*_0_ *be the multiplication of the ratios, such that for each (directed) edge* (*a, b*) *of the tree spanning S*_0_ *rooted at i*_0_, *the ratio R*_*a,b*_ *that goes from the genome at node a to the genome at node b is included in the multiplication;*

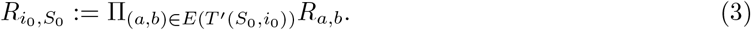

**Comment:** We are using this result later for *S*_0_, which consists of at most three leaves.

Note that for any two different nodes of the tree, 𝒢_*a*_ and 𝒢_*b*_, the *path ratio R*_*a,b*_ = *R*_*a,{a,b}*_ of the path 𝒢_*a,b*_ is a special case of a spread ratio. From the next theorem it follows that if *S*_0_ is a set of leaves, then 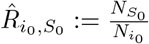 is an estimator for 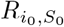.

#### Theorem 1.

*For a set of nodes S*_0_ *and a node i*_0_ ∈ *S*_0_, *if N*_0_ → ∞, *then* 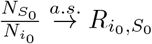.

The proof of the theorem, appearing in Subsection A.4 in the appendix, relies on the Convexity Lemma 1, ensuring that if a gene exists at the intersection of two nodes, then it exists along the entire path connecting them. For an example of a spread ratio, see Fig. 1d. The spread equation is explained in the next subsection.

### 2.5 The Spread Equation

Instead of working with multiplications like in Def. 1 and Thm. 1, it is possible to define the spread distance 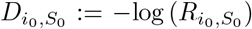 and denote its estimator by 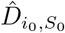 then obtaining the spread equation from *i*_0_ to *S*_0_;

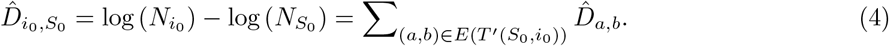

For every *S*_0_, which is a set of leaves, we obtain an equation such that the edge directional distances 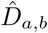 are the unknown of the equation (for an example, see Fig. 1d). We will need the following basic properties later.

**Properties**

a. **(Additivity)** If *a, b, c* are nodes such that *b* is on the path from *a* to *c*, then,

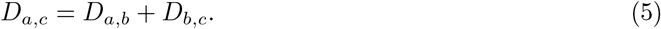
b. Let *a* and *b* be nodes and *N*_0_ → ∞ then

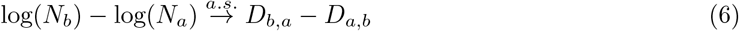 The last property connects the sizes of the nodes and the directional distances, and is applied later to obtain estimations for the tree nodes. Subsection A.5 is an extended version of this subsection.

### 2.6 Trees with Three Leaves

For a rooted tree with nodes 𝒢_0_, 𝒢_1_, 𝒢_2_, 𝒢_3_, 𝒢_4_, the root is 𝒢_0_ and the leaves are 𝒢_2_, 𝒢_3_, 𝒢_4_. The corresponding unrooted tree has nodes 𝒢_1_, 𝒢_2_, 𝒢_3_, 𝒢_4_. In Fig. 1e, we show the data items that we use. In Fig 1f, one can see the estimators of the edge directed distances we want to obtain. Each spread distance, 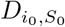, where *S*_0_ is a subset of leaves, provides an equation. There are nine possible spread equations for; *D*_2,3_, *D*_3,2_, *D*_2,4_, *D*_4,2_, *D*_3,4_, *D*_4,3_, *D*_2,*{*2,3,4*}*_, *D*_3,*{*2,3,4*}*_ and *D*_4,*{*2,3,4*}*_. We show that there is a unique solution for these equations. For a detailed extended version of this subsection, see subsection A.8.

## 3 The Triplet Directed Distance based Tree Reconstruction Algorithm

(For extended version of this section see Sect. B)

We now describe our cherry-peaking triplet-directed distance reconstruction (TDDR) algorithm in detail. The formal pseudocode description appears in Alg. 3.6. The previous section deals with a tree with three leaves. Therefore, we assume that we are dealing here with a tree with more than three leaves. Here, the original rooted tree is denoted by *T*. As explained in the appendix in Subsection A.7 it is only possible to reconstruct the unrooted related tree *T*_*u*_.

### 3.1 Reconstructing the Unrooted Tree for Every Triplet of Leaves

The TDDR algorithm begins by reconstructing an unrooted tree for each triplet of leaf genomes and estimating all directional distances. For every set of three distinct leaves *S* = *{i*_1_, *i*_2_, *i*_3_*}* there is a unique unknown internal node 𝒢_*c*_ with an index of *c* = *c*(*S*), which is the only node that exists on the path between any two members of *S*. We refer to *c* as the *center* of *S*. For any such triplet *S*, we can estimate directional distances from the leaves to the center 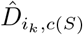 and from the center to the leaves 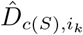, for *k* = 1, 2, 3 (see subsection A.8).

### 3.2 Estimating Directed Lengths of Edges Connected to Leaves

We now present an estimator for the directed distances of the leaf edges in *T*_*u*_. For any leaf 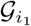, let 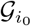 be the node in *T*_*u*_ that has a common edge with 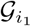. Denote 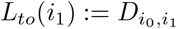 and 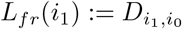.

To estimate *L*_*to*_(*i*_1_) (and similarly for *L*_*fr*_(*i*_1_)), we rely on the following observation: Let *U* (*i*_1_) be all triplets of leaves *S* such that *i*_1_ ∈ *S* and *m* is the number of leaves. Then, the *m −* 2 shorter distances of the form 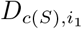 where *S* ∈ *U* (*i*_1_) are equal to *L*_*to*_(*i*_1_).

This is the reason why our estimator 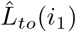 for *L*_*to*_(*i*_1_) is the mean of the shorter *m −* 2 estimates of distances of type 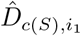 where *S* ∈ *U* (*i*_1_).

Similarly, the estimator 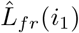 for *L*_*fr*_(*i*_1_) is the mean of *m −* 2 shorter estimates of distances of type 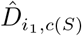 where *S* ∈ *U* (*i*_1_). For a formal definition of 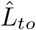 and 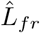, the proof of observation, and the proof of the consistency of the estimators, see subsection B.2 in the appendix.

### 3.3 Locating a Cherry

Extension of this part, including proofs and formal arguments, appears in subsection B.3 in the appendix. The algorithm presented is of cherry-picking type, similar to the neighbor-joining algorithm [74] in that a pair of leaves *a, b*, supposed to be a cherry, is sought in *T*_*u*_. Then, these two leaf sisters are removed from *T*_*u*_. By doing so, in the new tree 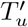, a subtree of *T*_*u*_, the sister’s parent node becomes a leaf, and the total number of leaves is reduced by one. We show how to repeat this procedure, locating cherries and removing their leaves, continuing to reduce the number of leaves, and, while doing so, assuming that the cherry-picking is correct, the original topology of *T*_*u*_ is reconstructed.

Here, we present our method for choosing such a cherry based on the estimators of directional distances for triplets of leaf genomes. Let *i*_1_ be a leaf, *i*_0_ be the (internal) node neighboring *i*_1_ and let *i*_2_ be another leaf. For the task of detecting a cherry pair, we define by *H*_*to*_ (*i*_1_, *{i*_1_, *i*_2_*}*) the mean of the distances from the centers of the triplets to *i*_1_ for all triplets *S* such that *i*_1_, *i*_2_ ∈ *S*. Similarly, *H*_*fr*_ (*i*_1_, *{i*_1_, *i*_2_*}*) is the mean distance from *i*_1_ to the same centers. We use the notation *Ĥ* to denote the means of the estimators of the correspondent distances.

We define 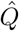 by subtracting the function 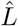 from *Ĥ* :

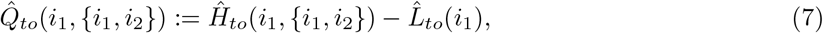

and

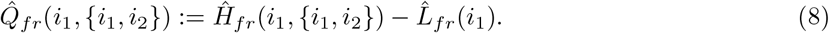

The functions 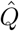 have the following desired properties: they are nonnegative, converge a.s. and converge to zero if and only if the pair *i*_1_,*i*_2_ form a cherry. Each pair of leaves *i*_1_ and *i*_2_ is graded using the four 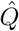 values;

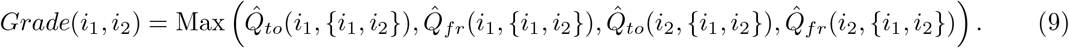

The pair of leaves *a, b* that is chosen as a cherry in the reconstructed tree minimizes the function *Grade*.

Let *c* be the node in *T*_*u*_ connected to *a* and *b*. By Eq. 6 We estimate *N*_*c*_ by geometric mean of two estimations:

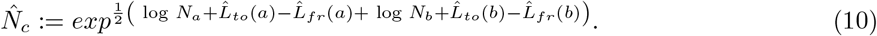

For the proof of statistical consistency and the last equation, see B.3.

### 3.4 Reducing the Number of Leaves by One

Denote by *i*_0_, *i*_1_ the indexes of the detected cherry sisters. Let us denote by *i*_2_ their parent index. Let 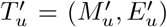 be the subtree of *T*_*u*_ after removing the sister leaves. To continue with 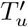 the same as we did with *T*_*u*_, we first need to estimate the directed distances of the triplets of leaves with *i*_2_ as one of the leaves. It was assumed that *T*_*u*_ has four or more leaves, so let *i*_3_ and *i*_4_ be any two other leaves of the tree *T*_*u*_. The directed distances of the triplet *{i*_2_, *i*_3_, *i*_4_*}* from 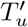 are estimated based on the triplets *{i*_0_, *i*_3_, *i*_4_*}, {i*_1_, *i*_3_, *i*_4_*}, {i*_0_, *i*_1_, *i*_3_*}, {i*_0_, *i*_1_, *i*_4_*}* and the directed distances of the leaf edges of *i*_0_ and *i*_1_ from *T*_*u*_. These estimators are calculated where *i*_3_ and *i*_4_ run over all pairs of leaves without *i*_0_ or *i*_1_. This gives all the estimators needed to pick the next cherry in 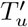. For a complete description and proofs, see Subsection B.4

### 3.5 The End of the Algorithm

At the end of the loop, there is only one triplet with three leaves, the directed distances from the center to the leaves of the triplet and vice versa. The center of the triplet is the last node that is added to the reconstructed tree, and if the triplet is *a, b, c* and the center is *d* then by Eq. 6 we estimate *N*_*d*_ by the geometric mean of three estimations:

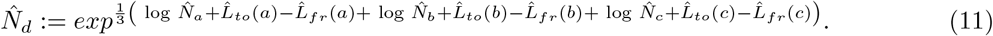

### 3.6 The Formal TDDR Algorithm

The following is the pseudocode of the algorithm:

#### Algorithm 1

TDDR

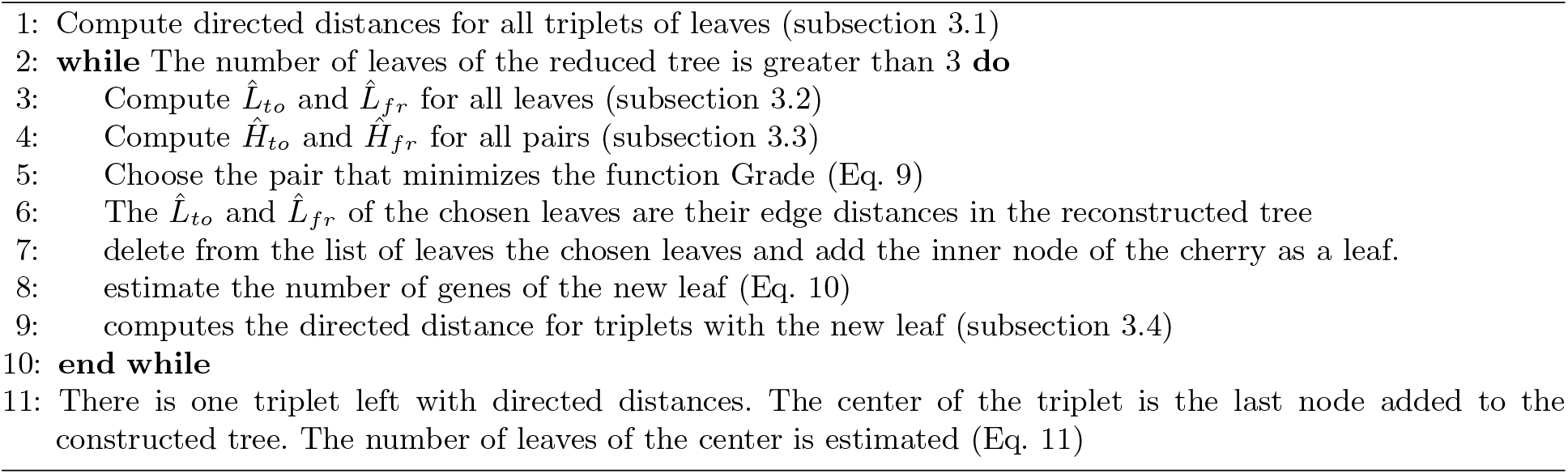

Assume that the data consist of a family of *m* leaf genomes and *n* is the maximum number of genes per genome, then the complexity of the first part (step 1) is *O*(*m*^3^*n*) due to the intersections computed for each genome triplet. The complexity of the second part (steps 2-11) is *O*(*m*^4^ log *m*) which is due to the computation of 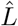 which consists of sorting a list of 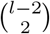 items for each of *l* leaves, where *l* goes from *m* to 3. Therefore, the complexity of the algorithm is *O*(*m*^3^*n* + *m*^4^ log *m*).

In Sec. B.7 of the appendix, we provide a toy example of reconstructing a tree from simulated data generated from a model tree over four leaves. The example demonstrates all the stages and calculations performed during the reconstruction.

## 4 Experimental Results

(For extended version of this section see Sect. C)

### 4.1 Simulation Study

Our experimental results start with a simulation study in which we tested the TDDR algorithm on synthetic data that we produced and to which we applied both the TDDR algorithm and two other popular packages - UNIMOG [10] and the Symmetric Gene Content employed by [78]. Two levels of experiments were run: simulation of a triplet of genomes and simulation over an entire tree, where all three approaches were tested, and the goal was to reconstruct as accurate as possible the model tree.

#### Simulations of Trees with Three Leaves

For simulations of trees with three leaves, we start with 𝒢_0_ with a size of 2000 genes, then produce from the parent 𝒢_0_ by two independent TRPs the children’s genomes 𝒢_1_ and 𝒢_2_ and then from 𝒢_1_ as parents we produce again by two independent TRPs the children 𝒢_3_ and 𝒢_4_. We perform simulations for mean directional edge lengths 0.005, 0.01, 0.015, …, 1; 10, 000 repetitions for each length. The 0.05 and 0.95 quantiles of the produced genomes, *N*_1_, *N*_2_, *N*_3_, and *N*_4_ are presented in Fig. 2a. We estimate the directional distances for the estimators presented in subsection A.8. The standard errors for estimators of the directed distances are shown in Fig. 2b. The errors for the different edges are close enough to justify the presentation of the combined error. We also estimate the size of the internal node 𝒢_1_ for the estimator presented in subsection A.8 and the standard errors are shown in Figure 2c.

**Fig. 2.**
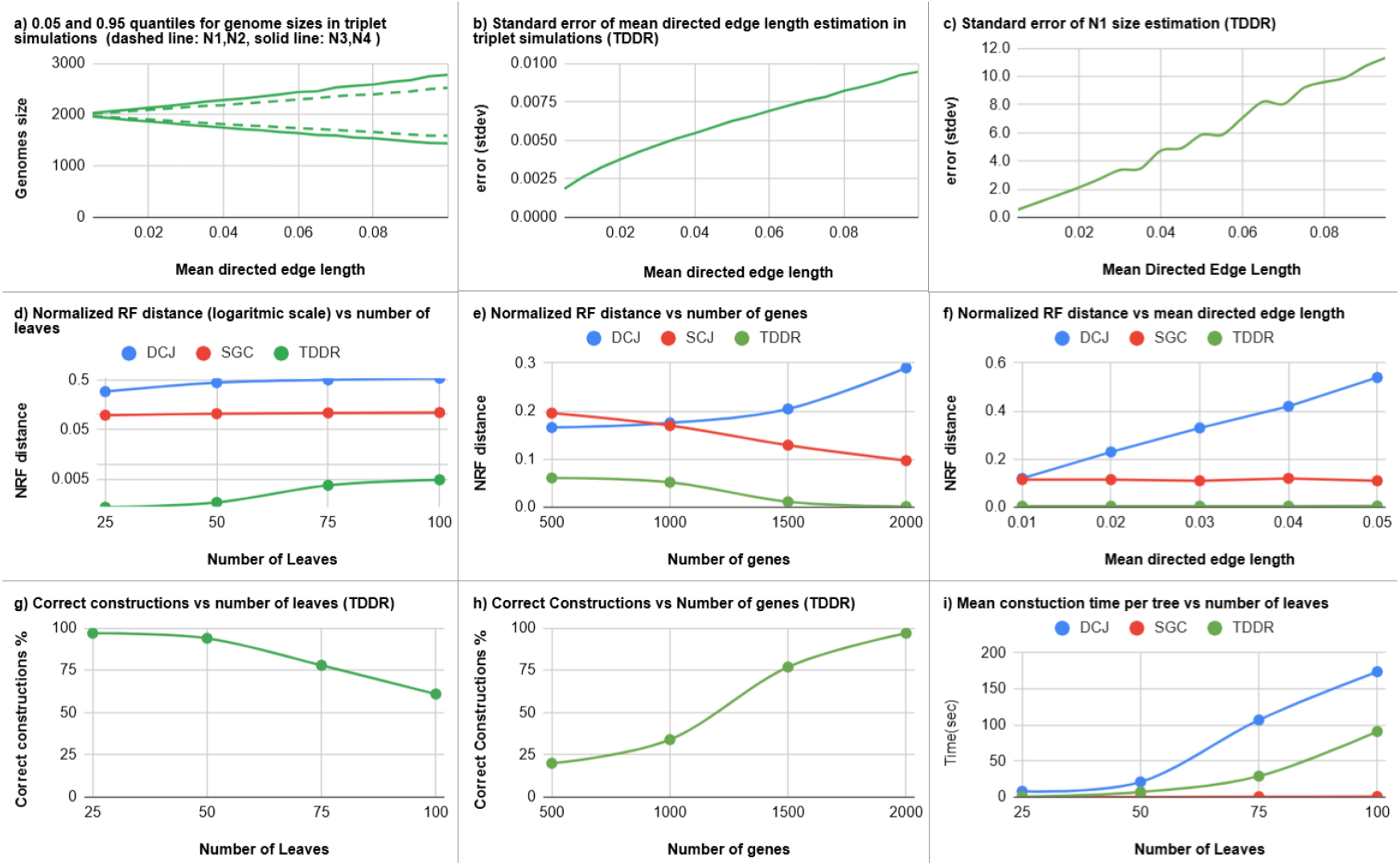
a) Sizes of genomes in simulations of trees with three leaves: for 10,000 repetitions and mean directed edge lengths 0.005 *−*0.1, these are the 0.5 and 0.95 quantiles for *N*_1_ and *N*_2_ and for *N*_3_ and *N*_4_. Genome sizes *N*_1_ and *N*_2_ have the same distributions, and so do *N*_3_ and *N*_4_. **b) Estimation error for the directed distances in simulations of trees with three leaves:** for 10,000 repetitions and mean directed edge lengths 0.005 *−*0.1, standard errors for the directed distances *D*_1,2_, *D*_2,1_, *D*_1,3_, *D*_3,1_, *D*_1,4_ and *D*_4,1_. **c) The estimation standard error for** *N*_1_ **in simulations of trees with three leaves:** the size of 𝒢_1_ for 10,000 repetitions and mean directed edge lengths 0.005 *−*0.1. **d) Normalized RF distance (logarithmic scale) vs number of leaves**. Mean directed edge length 0.05, Tree sizes: 25, 50, 75, and 100. For each size, 100 repetitions. **e) Normalized RF distance vs number of genes (size of** *N*_0_ ). Mean directed edge length 0.05, Tree size is 25 leaves, sizes of 𝒢_0_ : 500, 1000, 1500, and 2000. For each size, 100 repetitions. **f) Normalized RF distance vs mean edge length:** Tree size 100 leaves, mean directed edge lengths 0.01-0.05. for each mean length, 100 repetitions. **g) Percentage of correct reconstructed trees vs number of leaves**. Mean directed edge length 0.05, Tree sizes: 25, 50, 75, and 100. For each size, 100 repetitions. **h) Percentage of correct reconstructed trees vs number of genes**. Mean directed edge length 0.05, Tree size is 25 leaves. For each size, 100 repetitions. **i) Mean CPU time for one repetition vs number of leaves**. Mean directed edge length 0.05. Tree sizes: 25, 50, 75, and 100. For each size, 100 repetitions.

**Fig. 3.**
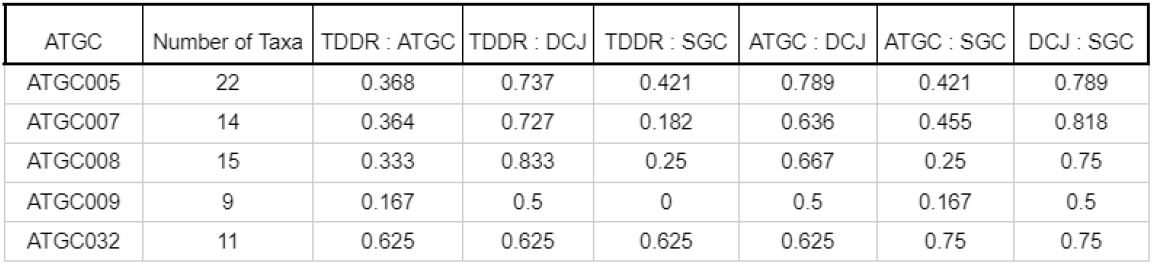
Normalized RF comparisons: The trees created by the three methods TDDR, DJC, SJC, and the tree downloaded from the ATGCs site are compared by normalized RF distance for the ATGCs: ATGC005, ATGC007, ATGC008, ATGC009, and ATGC032.

#### Simulations for the Tree Reconstruction Algorithm

The algorithm (TDDR) is described in Sect. 3. In each simulation repetition, a new (random) rooted tree is drawn starting with the genome *G*_0_, which has *N*_0_ genes at the root. Genome evolution proceeds recursively, starting from the root genome: A random leaf is chosen from the tree and two new child leaves are connected. Two independent TRPs create the genomes of these children of the new parent. This process continues until the desired number of leaves is reached. The leaf genomes resulting from this process are passed as input to the various packages under test. The TDDR is compared here with two different methods to create distance matrices implemented in the following packages;

a. Double-Cut-and-Join (DCJ) distance [8] (with indels) implemented by UNIMOG [10].
b. Symmetric Gene Content.

In both methods, the trees are reconstructed from the distance matrices by the neighbor-joining implementation of the BioPython package. Three sequences of simulations are made.

Seq. 1 The trees have sizes of 25, 50, 75, and 100 leaves with a mean directed edge length of 0.05. For each tree size, 100 repetitions.

Seq. 2 Mean directed edge lengths 0.01, 0.02, 0.03, 0.04, 0.05 with trees of size 100 leaves. For each mean length of the edge, 100 repetitions.

Seq. 3 First genome *G*_0_ with sizes (*N*_0_) 500, 100, 1500, 2000 with trees of size 25 leaves. For each genome size, 100 repetitions.

The reconstructed trees are tested against the true tree for all methods using the Robinson-Foulds distance implemented in the ETE3 package (Fig. 2d-f). We also tested for TDDR the correct reconstruction rates for Seq.1 and Seq.3 (Fig. 2g-h) and the mean time for tree reconstruction for the three methods (Fig. 2i).

To conclude this part, as a preface for the real data analysis, we can see that the TDDR algorithm performs reasonably well first in reconstructing edge lengths of the triplet and, even more impressively, by reconstructing the tree topology in the third part, in particular, compared to the other two packages that were tested.

### 4.2 Real Data Study - the Alignable Tight Genomic Clusters

The Alignable Tight Genomic Clusters (ATGCs) database [39] contains several million proteins from thousands of genomes organized into hundreds of clusters of closely related bacterial and archaeal genomes. Each ATGC is a taxonomy-independent cluster of two or more completely sequenced genomes. The ATGC database also includes many ATGC-COGs (Clusters of Orthologous Genes) and phylogenetic trees of the organisms within each ATGC. For the narrow phylogenetic spectrum of each ATGC, the probability of a duplication event inside an ATGC is very small [40,73]. In addition, orthology detection is significantly simplified compared to more general DB (e.g. EggNOG [29]). These facts make our model assumptions of infinite pangenome and a single copy COG highly realistic. We also note that no discordance between a gene-tree and the species-tree is possible under such a model. For the real data test, we used the following ATGCs; ATGC005, ATGC007, ATGC008, ATGC009, and ATGC032. The genome here is represented in SGC and TDDR as a set of distinct COGs (instead of genes). We used the same methods used for the simulations for tree reconstructions. We also used the phylogenetic tree downloaded from the ATGCs site. The various trees are compared by normalized RF and are given in Table 3. To conclude this part, we note that the differences between the directed distances on the same edge and between edges reflect real differences and support the need for the TRP model, untying gain and loss rates.

## 5 Conclusions

With the unprecedented availability of genomic sequences, the focus has shifted to large-scale reconstruction of both the phylogenetic density and the spectrum. Consequently, genome-wide approaches that leverage stronger signals than traditional phylogenetic markers have emerged. Among these approaches are genome dynamics (GD) techniques that capitalize on the relentless nature of such activity, especially in prokaryotes. However, these techniques have assumed relatively limited models for the complexity of modeling these events at high resolution.

In this work, we have introduced a new GD framework that models uncoordinated gene gain and loss rates. Our proposed extension is twofold: gain and loss rates may differ and change arbitrarily between edges (i.e., time and lineages) along the underlying evolutionary history. To our knowledge, this relaxation of the two rates has not been previously addressed, primarily due to the complications it imposes on modeling, particularly in phylogenetic reconstruction. An immediate implication of this extension is the loss of reversibility, a key concept in phylogenetics that overcomes the absence of time notion in contemporary molecular sequences. Another consequence is the relaxation of the constancy of genome size (in expectation). These consequences render standard phylogenetic approaches, particularly distance-based ones, futile.

Therefore, our second major contribution is an efficient model-based reconstruction algorithm, the triplet-based directed distances reconstruction (TDDR), which we prove to be statistically consistent. Our experimental work demonstrates that our algorithm can efficiently handle 100 taxa with high accuracy, significantly outperforming existing approaches not designed for data generated under this regime. Importantly, we applied TDDR to five datasets from the ATGC database, comparing it with two other approaches. Although our results show fair agreement with the sequence-based tree provided by ATGC, the more significant contribution of this experiment is its vivid demonstration of the proposed model’s necessity, as we observe outstanding differences in gain/loss rates along edges, refuting the adequacy of rate-equality models.

We acknowledge that components of this framework, either from the TRP model or the TDDR algorithm, have been used previously, such as varying rates of gene birth-and-death [33,5], non-stationarity and reversibility [81,11], or consistency analysis under several models [28,53,60]. While these components have been addressed in various past works under this or other applications, our complete framework - from the general model presented, through specializations to known models, to a dedicated algorithm based on bidirectional distances - represents a novel and interesting contribution.

Several future directions emerge from this work. On the algorithmic/theoretical side, we aim to refine our consistent result to guarantee reconstruction error for bounded genome size, particularly for real data sizes. We will work to improve the TDDR algorithm’s complexity to accommodate datasets of thousands of taxa, possibly using efficient sampling approaches with guaranteed marginal reconstruction error. On the practical side, we plan to apply TDDR to large ensembles of taxa sets to search for common mechanisms regarding gain/loss differences across genera and families.

## 6 Acknowledgment

The authors thank Eugene Koonin and Mike Steel for their helpful comments on the revised version.

## A The Model

(Extended version of Sect. 2)

In this part, we introduce the framework that accounts for different and changing rates of gain and loss and underlies this entire work. We describe a stochastic process in which a root genome evolves through binary-rooted phylogenetic tree to leaf genomes. Our building block is the two-ratio process (TRP) which represents a stochastic process that operates along the edges and paths of the tree *T*. Throughout this work, we assume that a genome is a set of unique genes (i.e., a single copy of every gene). A central underlying property used throughout the work is the *infinite pangenome model* [13,49], which implies that a given gene can be gained only once throughout history, maintaining the uniqueness of the gene in a genome. The following lemma, which we denote by *the convexity lemma* [45,44], follows directly from the infinite pangenome.

### Lemma 2.

It is also assumed that the TRP operates independently and individually along the directed edges.

A TRP denoted b y 𝒢_0,1_, has two parameters 0 *< R*_0,1_, *R*_1,0_ *<* 0 (the ratios). 𝒢_0,1_ starts with a parent genome 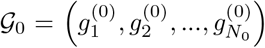, with *N*_0_ different genes, which evolves to a child 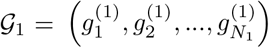 with *N*_1_ different genes. A gene in 𝒢_0_ *survives* to be in 𝒢_1_ with probability *R*_0,1_. Each gene 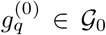 is replaced in 𝒢_1_ by a set *M*_*q*_ of genes, possibly including 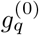 itself. All genes in *M*_*q*_ except 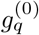 are new. Denote by *X*_q_ the number of genes in *M*. It is assumed that 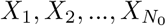 are i.i.d. with expectation *e*_0,1_ and variance *v*_0,1_. We define 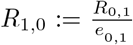. The variance is later considered to be a nuisance parameter. Fig. 4 illustrates the TRP on an edge. The number of genes that appear in both 𝒢_0_ and 𝒢_1_ is denoted by *N*_0,1_.

**Fig. 4.**
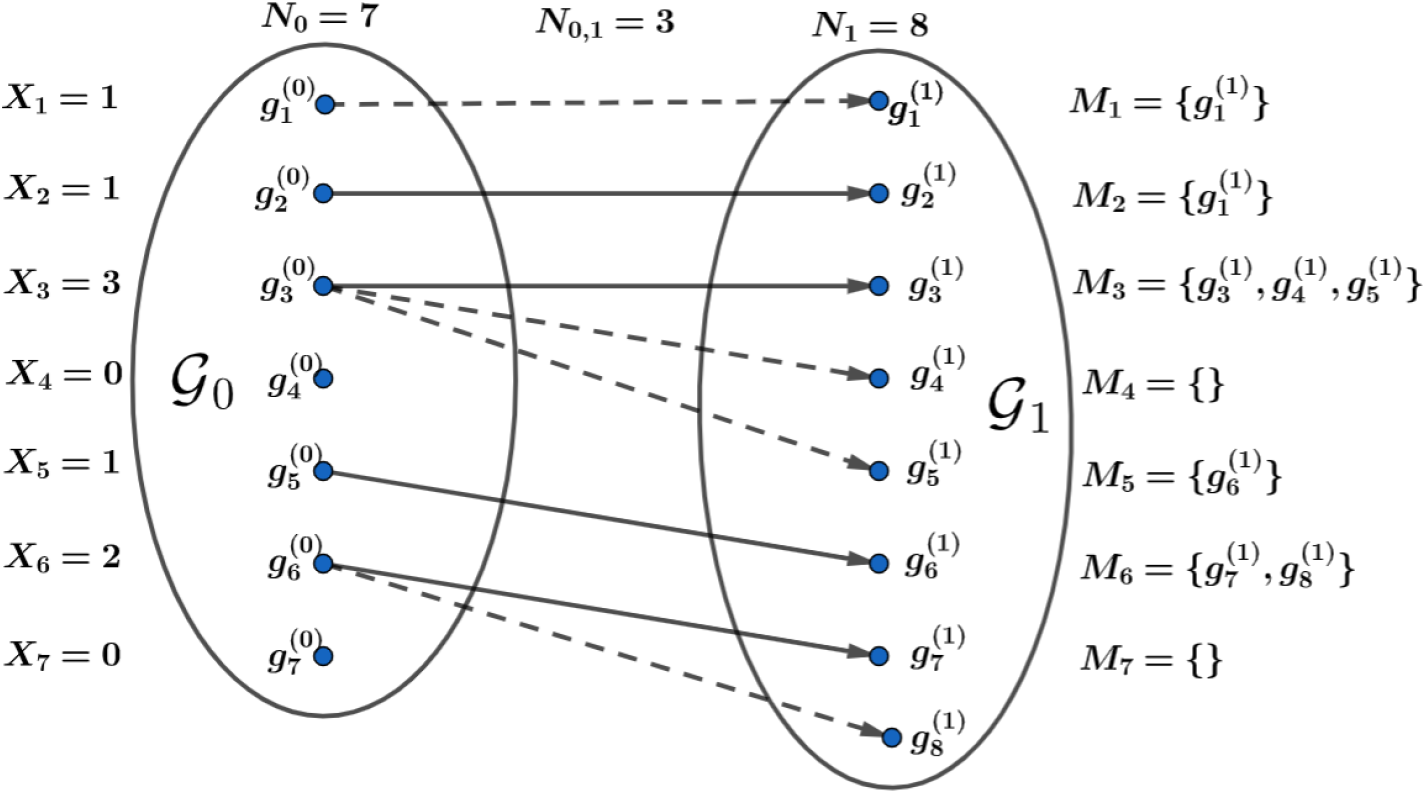
Two Ratio Process: The genome 𝒢_0_ evolves to genomes 𝒢_1_. The size of 𝒢_0_ is *N*_0_ = 7. The size of 𝒢_1_ is *N*_1_ = 8. Every gene 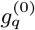 in 𝒢_0_ is replaced by a set *M*_*q*_ with a size greater or equal to zero. The gene 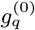 may appear 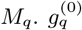 cannot appear *M*_*j*_, *j ≠ q*. An arrow goes from 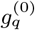 to the genes of the set *M*_*q*_ in 𝒢_1_. An arrow with the continuous line goes from a gene in 𝒢_0_ to the same gene in 𝒢_1_. The number of genes that appear both in 𝒢_0_ and in 𝒢_1_ is *N*_0,1_ = 3.

### Observation 3 (TRP Properties:)

*If N*_0_ *tends to infinity, then by the strong law of large numbers, we have*

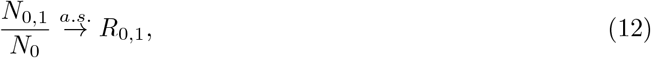

*Applying the strong law of large numbers, we again deduce that* 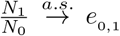. *Combining that with the previous item*,

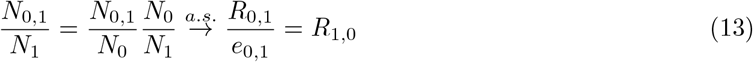

*In words, it means that* 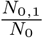 *and* 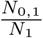 *tend almost surely to R*_0,1_ *and R*_1,0_ *respectively*.

From these properties, it follows that 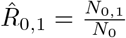 and 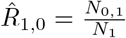 are estimators for the parameters *R*0,1 and *R*1,0.

**Comments:**

a. Although *R*_0,1_ is a probability, the ratio *R*_1,0_ is not a probability, and there is no reversed TRP process 𝒢1,0.
b. Later, when we deal with unrooted phylogenetic trees, the edges of the tree are TRPs where it is not known which of the two genomes starts and which ends the process. Since, at least in one direction, the ratio is not a probability, we prefer to refer to both as ratios.

### A.1 Birth-and-Death Process as a Special Case of the TRP

(Extended version of subsection. 2.1)

As mentioned above, the TRP model has, as a special case, several more popular models. Here, we show one. If we assume that 𝒢_0_ evolved to 𝒢_1_ by a linear birth-and-death process [20, p. 456] *n*_*t*_, 0 ≤ *t* ≤ *τ* where *τ* = *−*(log*R*_0,1_ + log*R*_1,0_), with a birth rate 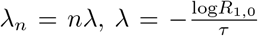 and a death rate *µ*_*n*_ = *nµ*, 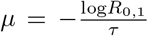, then it is possible to describe such a process between genomes as a TRP. The process begins with the population size *n*_0_ = *N*_0_, the size of 𝒢_0_. In time 0 ≤ *t* ≤ *τ*, the size of the genome is *n*_*t*_, then each gene can give birth to a new gene at a rate *n*_*t*_*λ*. It may also undergo a loss (death) event at a rate *n*_*t*_*µ*. Under these assumptions, the probability that a gene that was in 𝒢_0_ will also be after time *τ* in 𝒢_1_ is 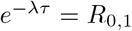. The expected number of genes in the set *M*_*q*_ in 𝒢_1_ that originated from a single gene 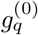 in 𝒢_0_ is 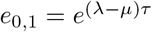, Then we have *e* 0,1 *R*0,1 = *R*_1,0_.

Nonhomogeneous birth-and-death, where *λ* = *λ*(*t*) and *µ* = *µ*(*t*) are functions of time, will also produce TRP. The sets *M*_*q*_ are defined similarly to the homogeneous case. Then 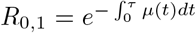, which is the probability that no event will occur in a nonhomogeneous Poisson process with intensity function *µ*(*t*)(see [54, p. 342]). For the computation of *R*_1,0_, one needs to know *e*_0,1_. To meet the TRP assumptions, the variance of the size of *M*_*q*_ must exist. For the expectancy and variance of the nonhomogeneous birth-and-death process, see [35].

### A.2 Composition of Two or More TRPs

(Extended version of subsection 2.2)

A process composed of two or more TRPs is a TRP(see Fig. 5). This property is essential for this work and is one of the advantages of the TRP model. It relies on the Convexity Lemma 1 defined above, guaranteeing that the genomes harboring a given gene form a connected subtree. For example, a composition of two or more birth-and-death processes with possibly different parameters is not necessarily a birth-and-death process. One can compose two nonhomogeneous birth-and-death processes, and the result is again a nonhomogeneous birth-and-death process, but the description of the TRP is more general and straightforward.

**Fig. 5.**
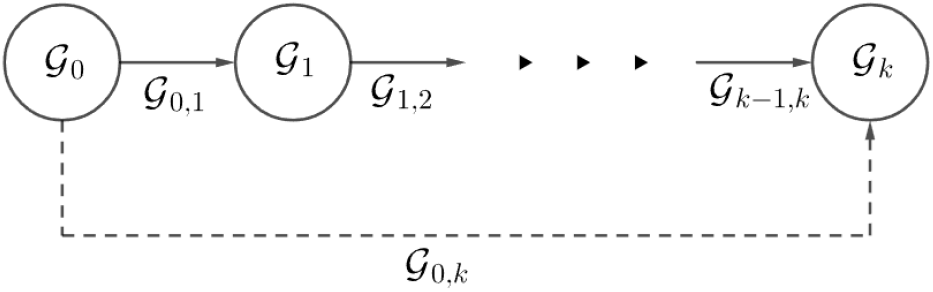
Composed Process of several TRPs: The genome 𝒢_*i*_ evolves to genomes 𝒢_*i*+1_ by the TRP 𝒢_*i,i*+1_. The composed process from 𝒢_0,*k*_ from 𝒢_0_ to 𝒢_*k*_ is also a TRP.

**Fig. 6.**
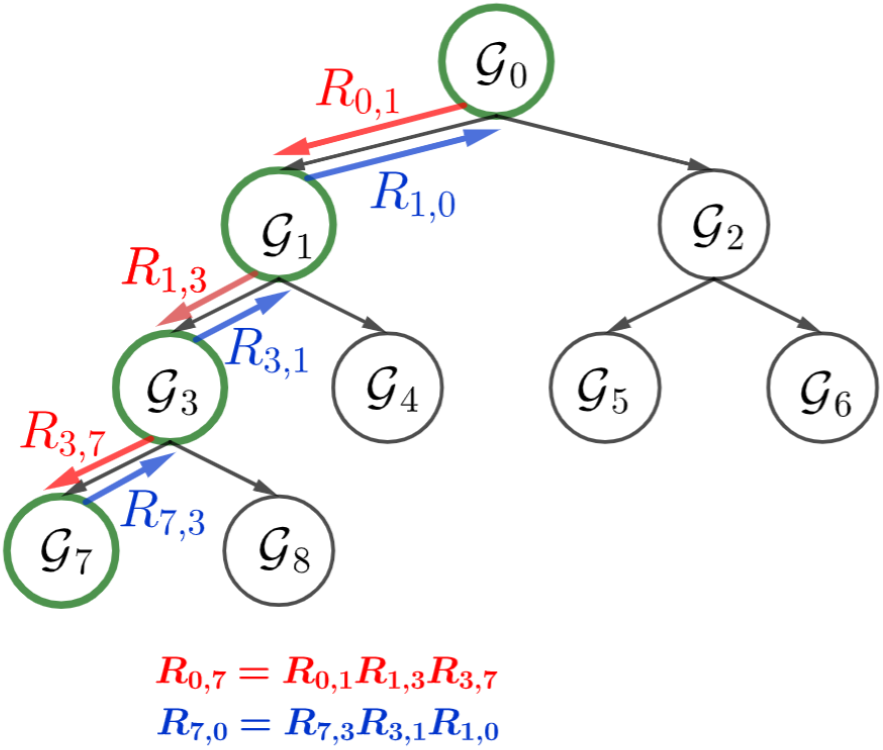
TRPs on a Phylogenetic Tree: There is TRP for each edge of the tree from parent to child. For example 𝒢_1,3_ from 𝒢_1_ to 𝒢_3_. Due to the composition the process 𝒢_0,7_ is also a TRP with ratios *R*_0,7_ and *R*_7,0_. 𝒢_1,7_ and 𝒢_0,3_ are also TRPs. Note the direction of the TRP is always down the tree. *G*_7,0_ is not a TRP.

Let 𝒢_0_, 𝒢_1_, …, 𝒢_*k*_ be a sequence of genomes such that 𝒢_*i*_ evolves to 𝒢_*i*+1_ by a TRP with ratios *R*_*i,i*+1_ and *R*_*i*+1,*i*_, and with expectation *e*_*i,i*+1_ and variance *v*_*i,i*+1_. By the Convexity Lemma 2, we have that a gene that underwent a loss event in one of the processes will not reappear later. We define that the gene 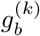 is in the set that replaced 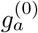 in 𝒢_*k*_, if there is a sequence of genes, 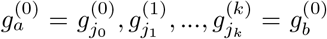, 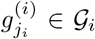, such that 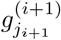 is in the sequence that replaced 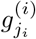 for 0 ≤ *i* ≤ *k −* 1. Then, the composed process 𝒢_0,*k*_ from 𝒢_0_ to 𝒢_*k*_ is also a TRP. The following settles the expectation and variance of the relations for the composition of *k* TRP and the process ratios.

#### Observation 4

*The composed process* 𝒢_0,*k*_ *is a TRP and*,

a. 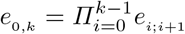
b. 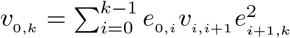 *(where e*= 1).
c. 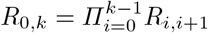
d. 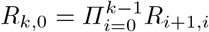

**Proof:** We prove this for *k* = 2. For *k >* 2, then the result follows by induction.

a. From the law of total expectation it follows that *e*_0,2_ = *e*_0,1_ *e*_1,2_.
b. From the law of total variance it follows that 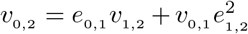. We need this only to show that the variance is finite to enable the use of the strong law of large numbers.
c. *R*_0,2_ is by definition the probability that a gene in 𝒢_0_ survives the two TRPs and reaches 𝒢_2_. This equality is true because of the Convexity Lemma 2.
d. Apply elements a, c and by definition 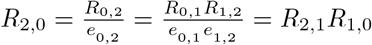.

### A.3 The Phylogenetic Tree

Let *T* = (*V, E*), where *V* = *{*𝒢_0_, 𝒢_1_, …, 𝒢_2*m−*2_*}*, be a connected rooted binary phylogenetic tree with *m* leaves. The convention here is that 𝒢_0_ is the root of the tree. The leaves represent known genomes because their sequence of genes is known, while the internal nodes represent unknown genomes. In the model, the information about the leaves that we consider includes the sizes of the leaf genomes and the intersection sizes among the members of pairs or triplets of the leaves.

Let *S* be a set of genome indexes. Denote the number of genes that appear in all these genomes by *N*_*S*_. We may omit parentheses of the set in the notation. Thus, we denote *N*_2,4_ instead of *N*_*{*2,4*}*_.

Then, if *i*_1_, *i*_2_, …, *i*_*m*_ are the indexes of the leaves, we use them as data for our estimations of the genome size of the node, 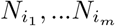, the intersections of pairs of genomes, 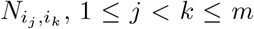, and the intersection of triplets of genomes, 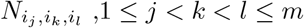.

For each directed edge, with neighboring node genomes 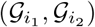, for which 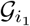 is considered a parent and 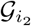 is its child, it is assumed that 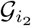 evolved from its parent through a process denoted by 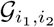.

This process acts according to the TRP model described in the previous section. Due to the composition property, even if *i*_2_ is just a descendant of *i*_1_, and not a direct child, then 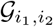 is also a TRP (see Fig. A.3). In this part, we describe a method for inferring the tree structure given leaf genomes assumed to have evolved via TRPs.

### A.4 Spread Ratio

(Extended version of subsection 2.4)

We first describe the path ratio, which is a special case of the spread ratio.

#### Path Ratio

Recall that for a pair of genomes 𝒢_*a*_ and 𝒢_*b*_ in *V*, the directed path 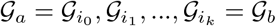. This path from 𝒢_*a*_ to 𝒢_*b*_ is indicated by 𝒢_*a,b*_ = (*i*_0_, *i*_1_, …, *i*_*k*_) (see example in Fig. 7). Note that the path is not necessarily a TRP.

**Fig. 7.**
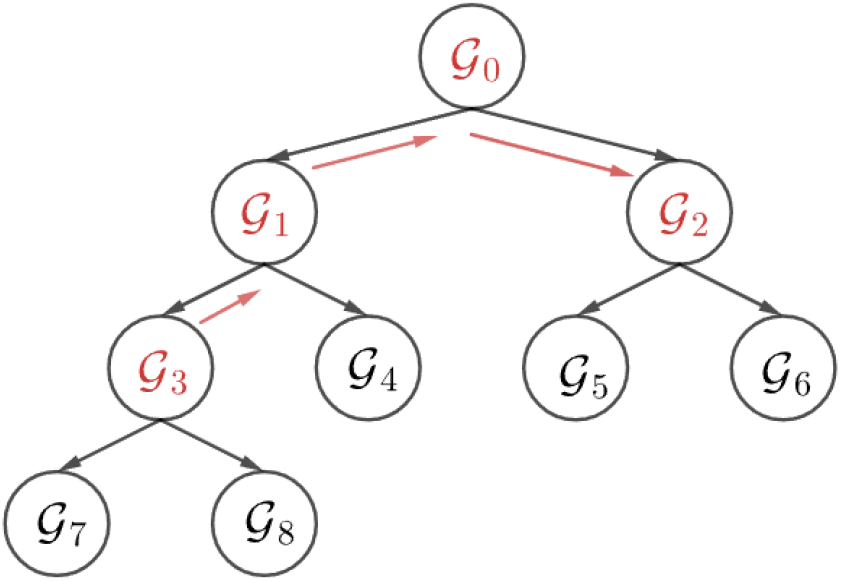
Path Ratio: The path ratio of the path 𝒢_3,2_ from 𝒢_3_ to 𝒢_2_ is defined as *R*_3,2_ = *R*_3,1_*R*_1,0_*R*_0,2_. 𝒢_3,2_ is not a TRP.

This assumption makes it possible to define the path ratio *R*_*a,b*_ for a path 𝒢_*a,b*_ between 𝒢_*a*_ to 𝒢_*b*_ by multiplying the proper of edges ratios;

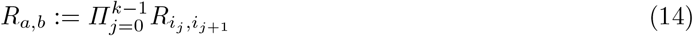

The following theorem justifies the definition of the path ratio;

##### Observation 5

*If N*_0_ → ∞ *then* 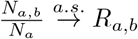

The proof of the following theorem will also cover the proof of the observation.

##### Corollary

If *N*_0_→ ∞, then by observation, for any nodes *a, b* that belong to the tree satisfy 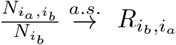 and 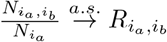. Dividing, we obtain

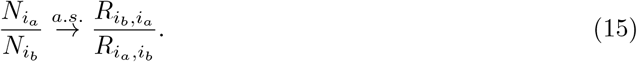

#### Spread Ratio

Given a set *S*_0_ of genome indexes at the nodes of *T*, let 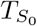 be the minimal connected subtree of *T* that contains *S*_0_, and let 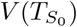 and 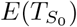 be the set of nodes and edges in 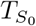, respectively. Now, for *i*_0_ ∈ *S*_0_, let *T*′(*S*_0_, *i*_0_) be the tree 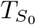 that is rooted at *i*_0_ ∈ *S*_0_, such that the (directed) edges (*a, b*) in *E*(*T*′(*S*_0_, ∈ *i*_*o*_)) are directed away from *i*_0_, i.e. node *a* is closer to root *i*_0_ than *b*.

##### Definition 2.

*For a set S*_0_ *of nodes in T and a node i*_0_ ∈ *S*_0_, *let the spread ratio* 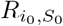 *from i*_0_ *to S*_0_ *be the multiplication of the ratios, such that for each (directed) edge* (*a, b*) *of the tree spanning S*_0_ *rooted at i*_0_, *we take in the multiplication of the ratio that goes from the genome at node a to the other genome of that edge;*

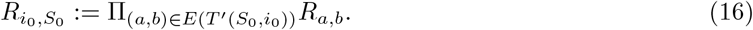

***Comment:*** *We are using this result later for S*_0_, *which consists of at most three leaves*.

Note that for any two different nodes of the tree, 𝒢_*a*_ and 𝒢_*b*_, the *path ratio R*_*a,b*_ = *R*_*a,{a,b}*_ of the path 𝒢_*a,b*_ is a special case of a spread ratio.

Fig. 8 illustrates the spread ratio in the leaf set *{*𝒢_4_, 𝒢_2_, 𝒢_7_, 𝒢_8_*}* with respect to rooting in the leaf *G*_4_. The next theorem is a generalization of Obs. 5. From this theorem it follows that if *S*_0_ is a set of leaves, then 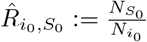 is an estimator for 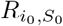.

**Fig. 8.**
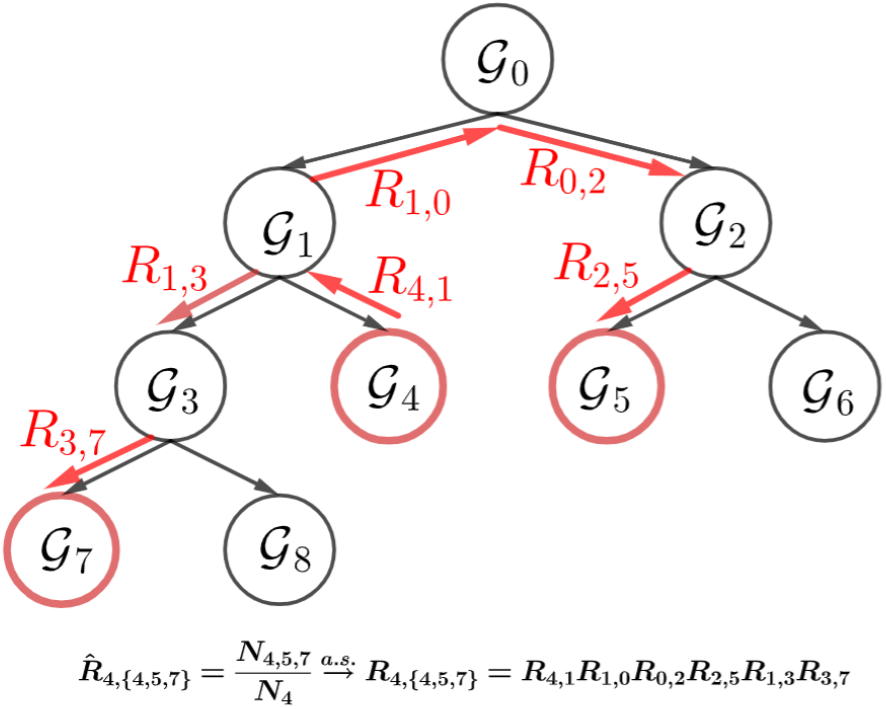
Spread ratio: The Spread ratio from 𝒢_4_ to *{*𝒢_5_, 𝒢_7_*}* is *R*_4,*{*4,5,7*}*_ = *R*_4,1_*R*_1,0_*R*_0,2_*R*_2,5_*R*_1,3_*R*_3,7_. The multiplication includes the ratios for directed edges of the tree rooted at 𝒢_4_ and has *{*𝒢_5_, 𝒢_7_*}* as its leaves.

##### Theorem 2.

*For a set of nodes S*_0_ *and a node i*_0_ ∈ *S*_0_, *if N*_0_→ ∞, *then* 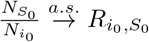.

The proof of the theorem is based on the Convexity Lemma 2, ensuring that sets of genes exist at the intersection of two leaves along the entire path connecting them.

**Proof:** As was previously mentioned in the assumption, all genomes possess a specific gene from a connected subgraph of the tree. Therefore, 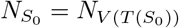 and it is enough to prove that 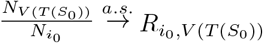 for any *i*_0_ in *V* (*T* (*S*_0_)). We prove it by induction on *i*_0_, starting with the case where *i*_0_ indicates the original root of *T* (*S*_0_) and then moving down the tree from parent to child. In the case where *i*_0_ is the original root of *T* (*S*_0_), all expression ratios are real probabilities, and the result of multiplication is the probability that a gene that is in the root of *V* (*T* (*S*_0_)) will also be in all genomes of *V* (*T* (*S*_0_)). Simple multiplication is possible due to the lack of memory in an edge process, and independence from all edges comes from preceding processes. This means that what happens to a gene in the process does not depend on its history from previous processes. Next, we show that if it is true for a parent node *i*_1_ in *V* (*T* (*S*_0_)), then it is also true for his child *i*_2_ in *V* (*T* (*S*_0_)). If it is true for the parent, then

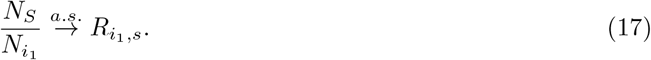

from Eq 15 it follows that

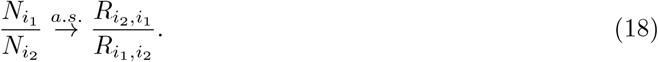

Thus,

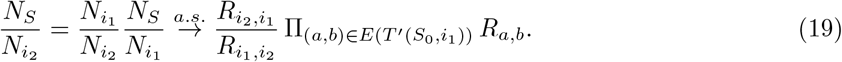

Note that *E*(*T*′(*S*_0_, *i*_1_)) and *E*(*T*′(*S*_0_, *i*_2_)) are almost the same, with the only difference that (*i*_1_, *i*_2_) ∈ *E*(*T*′(*S*_0_, *i*_1_)) and (*i*_2_, *i*_1_) ∈ *E*(*T*′(*S*_0_, *i*_2_)). It follows that

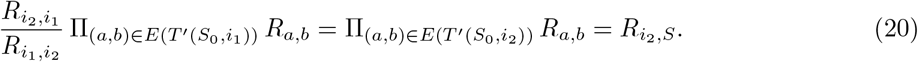

This completes the proof. ▪

### A.5 The Spread Equation

(Extended version of subsection 2.5)

Instead of working with multiplications like in Def. 2 and Thm. 2, it is possible to define directional edge distance 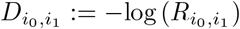 then define spread distance (and as a special case, path distance) from *i*_0_ to *S*_0_;

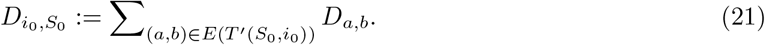

Thus, instead of multiplying probabilities or directional ratios along the edges, we use directional distances. Then we define the following estimator for 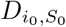;

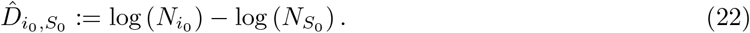

as a result of Thm. 2,

#### Observation 6

*If N*_0_ → ∞, *then*

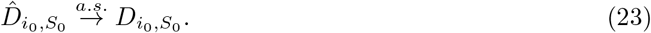

Combining Def. 2, 22 and this observation we obtain the spread equation (linear equation):

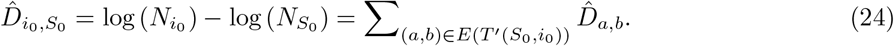

For every *S*_0_, which is a set of leaves, we obtain an equation such that the edge directional distances of the form 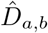 are the unknown of the equation (for an example, see Fig. 9).

**Fig. 9.**
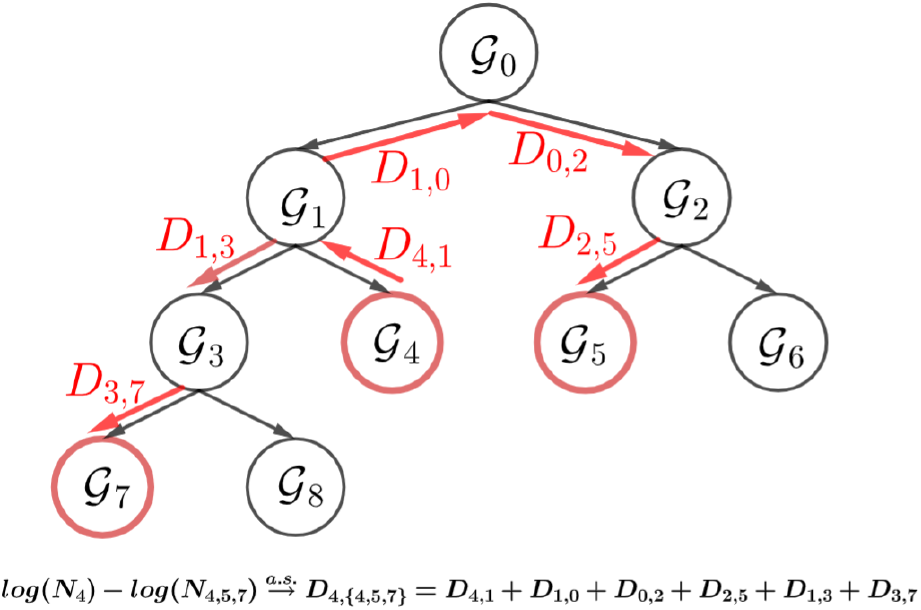
Spread equation: The spread ratio *R*_4,*{*4,5,7*}*_ from *G*_4_ to *G*_5_ and *G*_7_ is defined by the multiplication of the edge ratios that are notated by the red arrows. An estimator for this spread ratio is 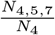.Applying the −log function, we obtain 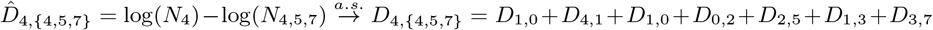. Thus, the spread equation is 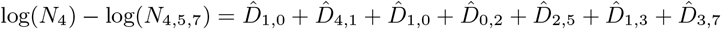.

We will need the following basic properties later.

#### Properties

a. **(Additivity)** From Eq. 21 we deduce that if *a, b, c* are nodes such that *b* is on the path from *a* to *c*, then,

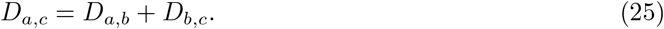
b. Let *a* and *b* be nodes and *N*_0_ → ∞ applying log to Eq. 15 we obtain that

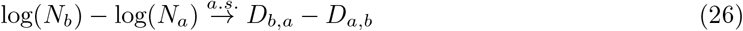 The last property relates between the sizes of the nodes and the directional distances. This is applied later to obtain estimations for the tree nodes.

### A.6 Reduction to Symmetric Distance

It is possible to define the following symmetric distance between two nodes of the tree: Let *a, b* be nodes of the tree *T* then define

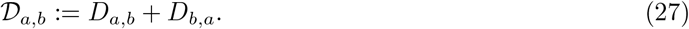

This distance is additive;

#### Observation 7

*Let a, b and c be nodes of T and assume that b is on the path between a and c. Then* 𝒟*a,c* = 𝒟*a,b* + 𝒟*b,c*.

**Proof:**

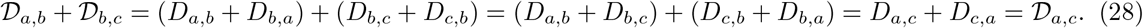

From Eq. 22 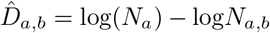 and 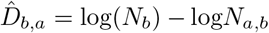 and therefore the natural estimator of the symmetric distance is

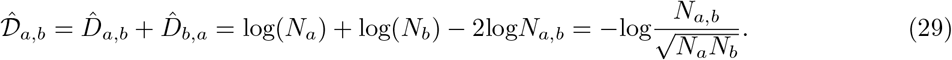

One can apply this to compute the distance matrix for the leaves and then use neighbor-joining to reconstruct a tree. However, this method uses less information compared to the method proposed here and, therefore, is less accurate in tree reconstruction.

### A.7 Infering the Unrooted Tree

The real original phylogeny is rooted like the tree on the right side of Fig. 10, and therefore also directed, with the edge directed away from the root toward the leaves. With the type of data at hand, that is, leaf genomes in the form of gene sets, the notion of time or edge direction is lost, and as a result, we can only infer the unrooted version of the original tree, such as the tree on the left side of the figure. This tree *T*_*u*_ = (*M*_*u*_, *E*_*u*_) is obtained by removing the root 𝒢_0_, placing an edge between its two child nodes and erasing the directionality of all edges. The directed path distances and spread distances on the unrooted trees are defined naturally by their correspondents in the rooted tree. We aim to obtain estimators for the edge directed distances of the unrooted tree. One of the edges of the unrooted tree represents a path of two edges connected to the root in the rooted tree. It is not possible for us to detect that the edge of the unrooted tree is the path of two edges on the rooted tree.

**Fig. 10.**
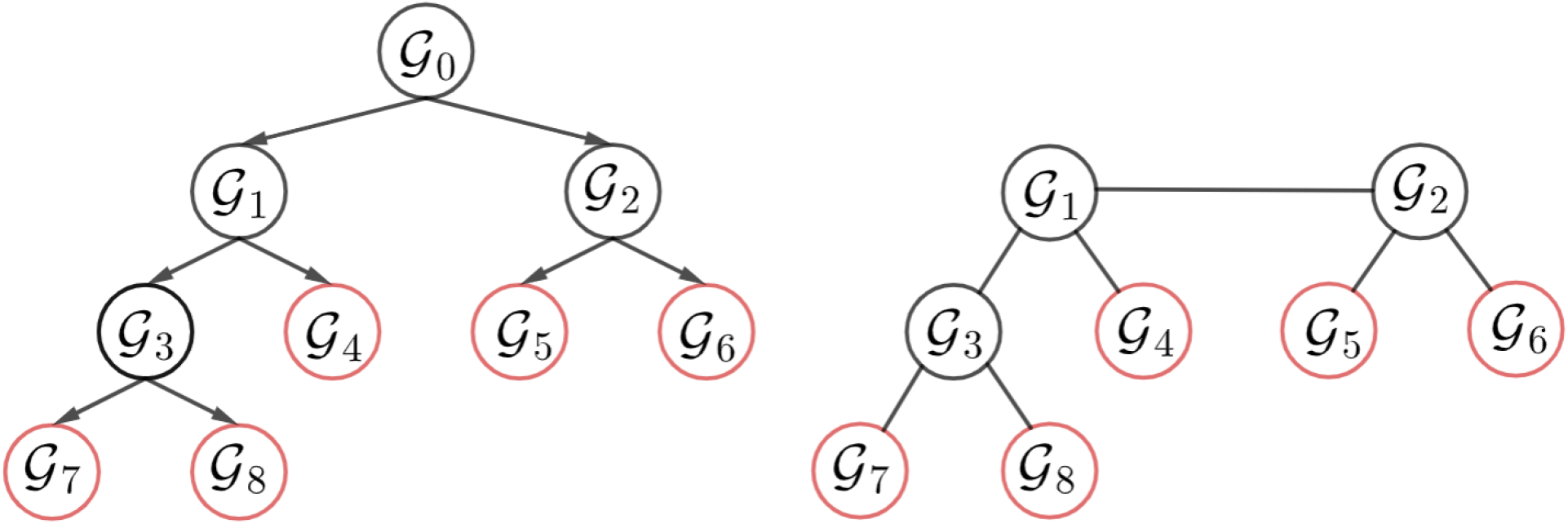
Rooted vs Unrooted: The original phylogenetic tree of the process, *T* = (*V, E*) is rooted like the tree on the left. With the type of methods we use, we can only infer an unrooted tree, such as the tree *T*_*u*_ = (*V*_*u*_, *E*_*u*_) on the right. The tree on the right originated from the tree on the left by erasing 𝒢_0_, placing an edge between 𝒢_1_ and 𝒢_2_, and erasing the polarization of the edges.

### A.8 Trees with Three Leaves

(Extended version of subsection 2.6)

We show that for a tree with three leaves, there is a unique solution that satisfies the spread equations of subsection A.5.

In Fig. 11, one can see the three possible trees with three leaves. It is possible to estimate all the path distances (or ratios) except the one for the directed paths that start or end with the root. In figure 12 on the left side we show the data items we use. On the right side, one can see the estimators of the edge-directional distances that we want to obtain. For each spread distance, 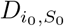 where *S*_0_ consists of leaves provides an equation. For a tree of three leaves 𝒢_2_, 𝒢_3_, 𝒢_4_ there are nine spread distances; *D*_2,3_, *D*_3,2_, *D*_2,4_, *D*_4,2_, *D*_3,4_, *D*_4,3_, *D*_2,*{*2,3,4*}*_, *D*_3,*{*2,3,4*}*_ and *D*_4,*{*2,3,4*}*_ and there are nine equations, respectively;

**Fig. 11.**
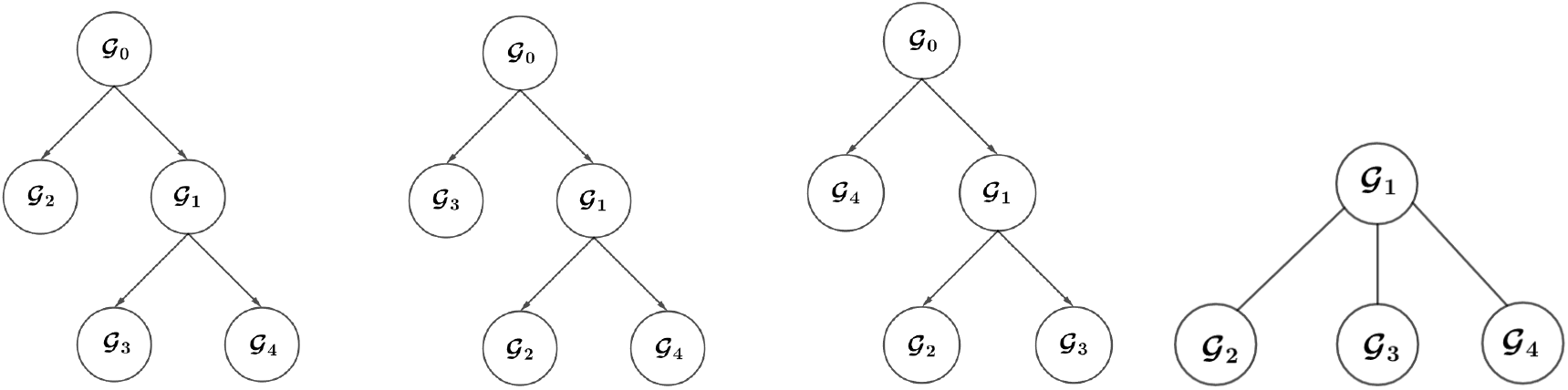
Trees with three leaves: The genome 𝒢_0_ is the root. The genomes 𝒢_0_ and 𝒢_1_ are internal nodes, 𝒢_2_, 𝒢_3_, and 𝒢_4_ are the leaves of the rooted binary tree and their sets of genes are known. There are three possible binary trees with roots. The three rooted trees produce the same unrooted tree on the right side. Observation 8 is true for all three trees.

**Fig. 12.**
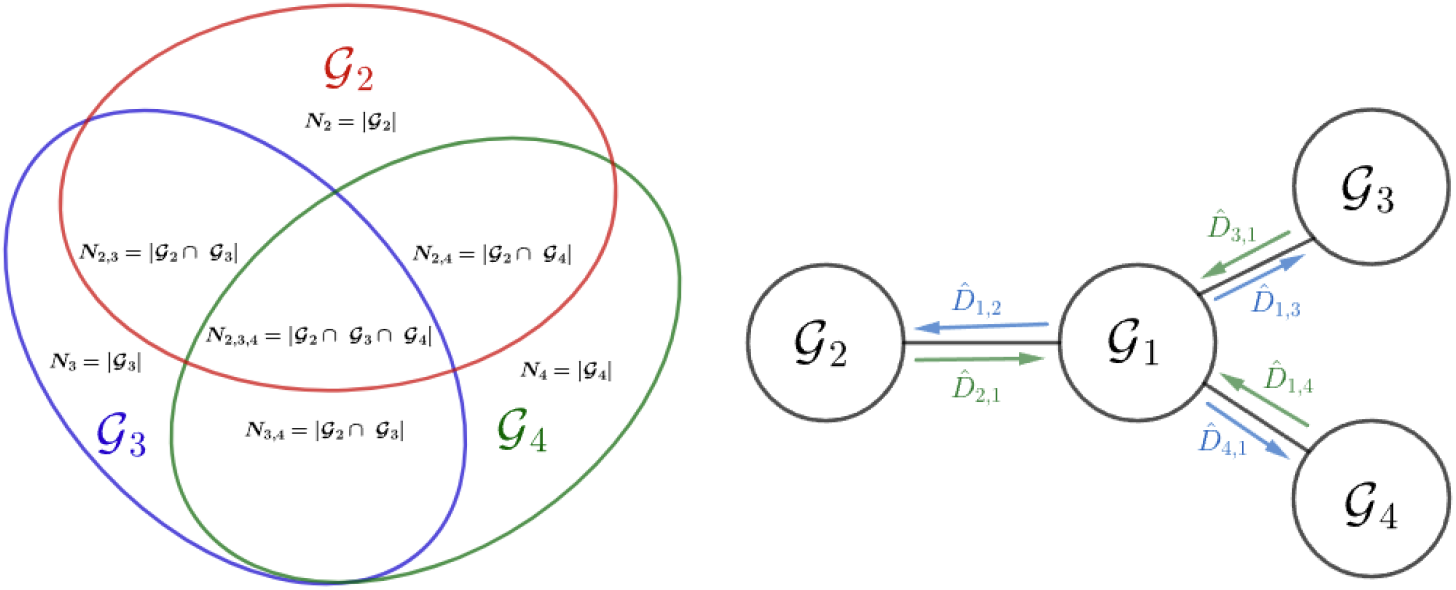
(L) The data of three leaves: the data of three leave consist of the genome sizes, the sizes of the intersections of pairs of genome and the size of intersection of all three genomes. **(R) The edge-directed distances are the unknowns:** the edge-directed distances from 𝒢_1_ to the leaves and the directed distances from the leaves to 𝒢_1_ are the unknown parameters that need to be estimated. There is a linear transformation with kernel of dimension 1 between the vector of logs of the sizes on the left and the vector of the unknowns that appears on the right. If the size of the intersection of all three is omitted, then there is not enough data to obtain the estimates.

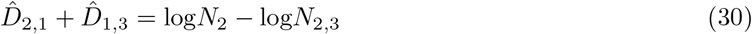

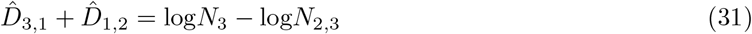

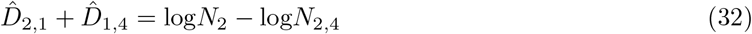

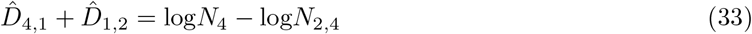

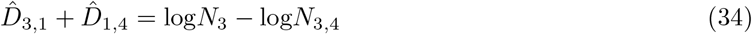

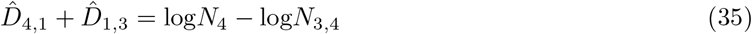

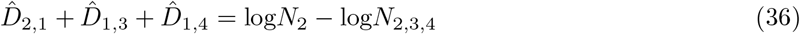

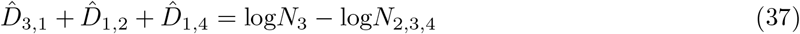

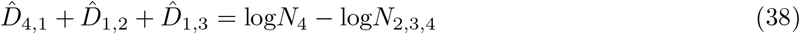

We show here that there is a unique solution that is given by a linear transformation between the vector of data items (log*N*_2_, log*N*_3_, log*N*_4_, log*N*_2,3_, log*N*_2,4_, log*N*_3,4_, log*N*_2,3,4_) and the vector of the unknowns consist of estimates of the edge-directed distances 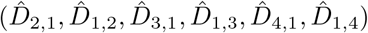. This transformation has a non-trivial kernel with dimension 1. If we add to all the components of the data vector a constant, the solution should remain the same. However, each of the data components is essential for the estimation. If we eliminate one of the data components, it is no longer possible to obtain estimates for all edges-directed distances. This justifies why later, for larger trees, we compute for each triplet of leaf genomes the size of their intersection.

#### Theorem 3.

*The following is the unique solution for equations 30-38*

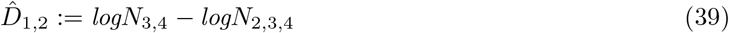

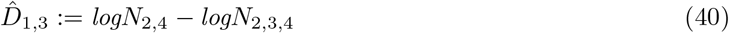

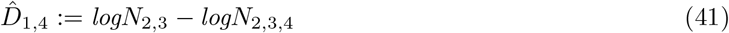

*and*

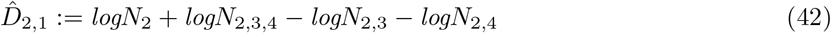

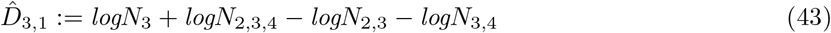

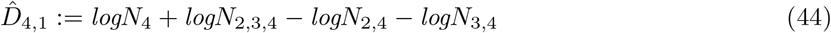

***Proof:*** *To show that this is solution, one needs to substitute the solution in the equations. Due to symmetry, it is enough to show this for equation 30 and equation 36 (one of each type). Substituting in equation 30:*

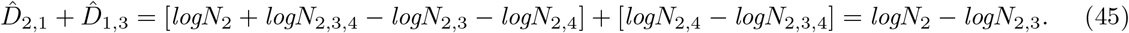

*Substituting in equation 36: In the previous equation, we calculated the sum of the first two components*,

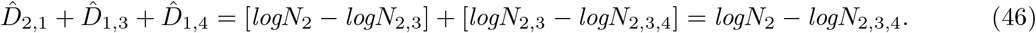

*To show the uniqueness of the solution, we need to derive the solution 39- 44 from equations 30-38*. *To get Eq 39 subtruct 35 from 38, for Eq 40 subtruct 33 from 38, and for Eq 41 subtruct 31 from 37*. *Since we obtained equations 39-41 we can use them to derive equations 42-44; for Eq 42 subtruct Eq 40 from Eq 30, for Eq 43 subtruct Eq 39 from Eq 31, and for Eq 44 subtruct Eq 39 from Eq 33*. ▪

In matrix notation this is;

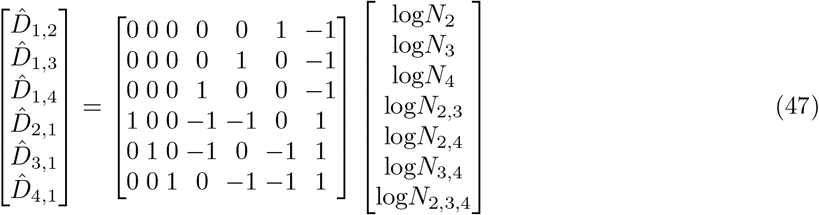

It can be seen on this matrix that the kernel of this transformation has a dimension of 1. Every data item is necessary. This is true in particular for the intersection of the three leaves *N*_2,3,4_, which is a relatively nontrivial data item. One cannot solve the equations for a tree with three leaves using only the sizes and the intersection of pairs.

#### Observation 8

*The estimators in the definitions 39- 44 are consistent*.

**Proof:** The proof of the consistency of the estimator 39 is as follows;

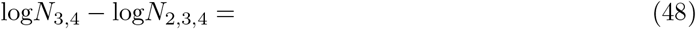

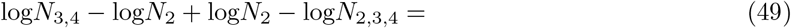

by definition 22 and observation 6,

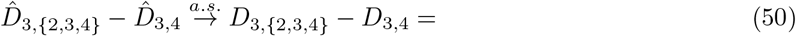

and by definition 21,

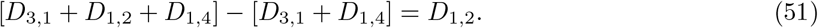

The consistency proofs for the estimators in the definitions 40-41 are similar. The proof of the consistency of the estimator 42 is as follows;

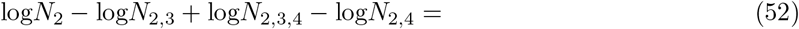

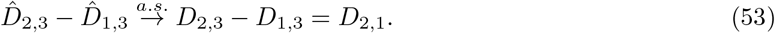

The consistency proofs for the estimators in the definitions 43-44 are similar. ▪

In addition, if we define,

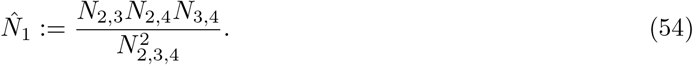

Then, as the next theorem shows, 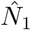 is an estimator for *N*_1_, which is an unknown random variable.

#### Theorem 4.

*If N*_0_ → ∞, *then* 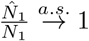.

**Proof:** From Theorem 8 we have

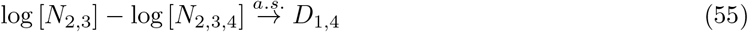

and

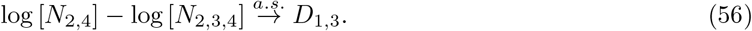

From observation 6 it follows that

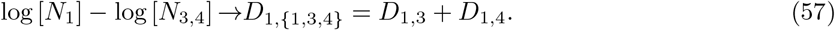

Subtracting expressions 55 and 56 from expression 57, we obtain

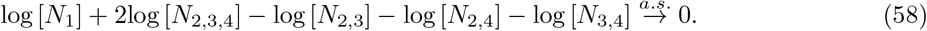

This is equivalent to

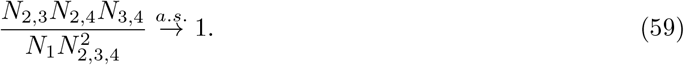

Estimation of *N*_1_ is a byproduct and is not essential for reconstructing the tree topology of larger trees.

## B The Triplet Directed Distance based Tree Reconstruction Algorithm

(Extended version of Sect. 3)

We now describe in detail our cherry-peaking triplet-directed distance reconstruction (TDDR) algorithm. The formal pseudocode description appears in Alg. B.6. The previous section deals with a tree with three leaves. Therefore, we assume that we are dealing here with a tree with more than three leaves. Here, the original rooted tree is denoted by *T*. As explained in the appendix in Subsection A.7 it is only possible to reconstruct the unrooted related tree *T*_*u*_.

### B.1 Reconstructing the Unrooted Tree for Every Triplet of Leaves

(Extended version of Sect. 3.1)

The TDDR algorithm begins with the reconstruction of an unrooted tree for each triplet of leaf genomes with the estimation of all directional distances, according to estimators 39-44. For every set of three distinct leaves *S* = {*i*_1_, *i*_2_, *i*_3_} there is a unique unknown internal node 𝒢_*c*_ with an index of *c* = *c*(*S*), which is the only node that exists on the path between any two members of *S*. We refer to *c* as the center of *S* (see Fig. 13). For any such triplet *S*, we can estimate directional distances, according to the estimators 39-44, from the leaves to the center, 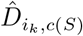 and from the center to the leaves, 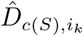, for *k* = 1, 2, 3.

**Fig. 13.**
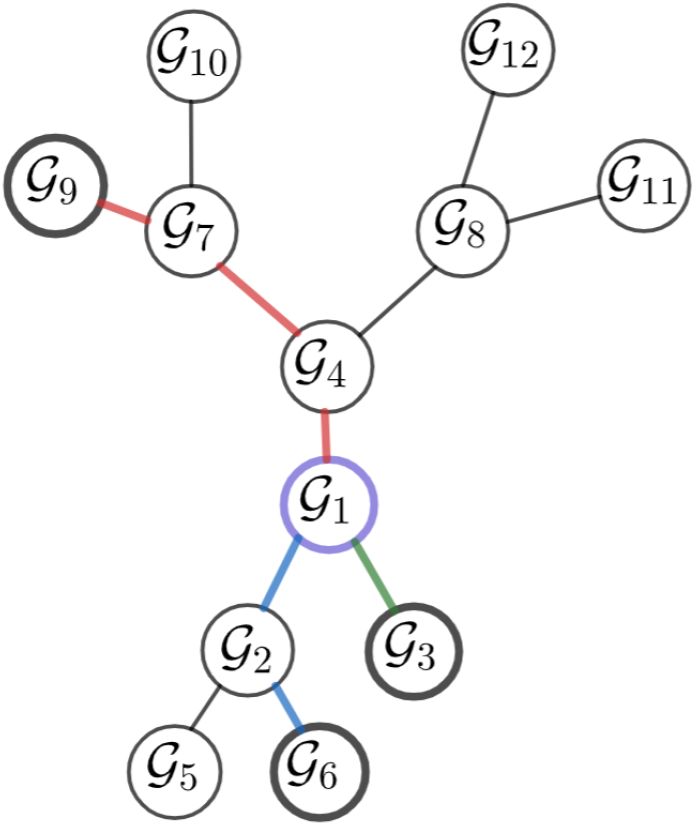
Every three leaf genomes have a unique center. The center for *{*𝒢_3_, 𝒢_6_, 𝒢_9_*}* is 𝒢_1_.

### B.2 Estimating Directional Lengths of Edges Connected to Leaves

(Extended version of Sect. 3.2)

We now present an estimator for the directed distances of the leaf edges in *T*_*u*_. For any leaf 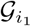, let 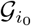 be the node in the unrooted tree that has a common edge with 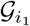. Denote 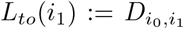 and 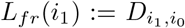. To estimate *L*_*to*_(*i*_1_) (and similarly for *L*_*fr*_(*i*_1_)), we rely on the following observation, which is explained later. Let *U* (*i*_1_) be all triplets of leaves *S* such that *i*_1_ ∈ *S* and *m* is the number of leaves. Then, the *m −* 2 shorter distances of the form 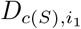 where *S* ∈ *U* (*i*_1_) are equal to *L*_*to*_(*i*_1_).

This is the reason why our estimator 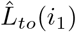 for *L*_*to*_(*i*_1_) is the mean of the shorter *m −* 2 estimates of distances of type 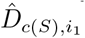 where *S* ∈ *U* (*i*_1_).

Similarly, the estimator 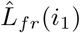 for *L*_*fr*_(*i*_1_) is the mean of *m −* 2 shorter estimates of distances of type 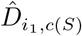 where *S* ∈ *U* (*i*_1_). The formal description of the observation and its proof

#### Observation 9

*There are at least m −* 2 *different triplets such that i*_1_ *is one of the leaves of the triplet and i*_0_ *is the center of the triplet*.

**Proof:** The node *i*_0_ divides the *m −* 1 leaves which are not *i*_1_ into two classes; two edges are in the same class if the path between them does not include *i*_1_ (for examples, see Fig. 14). Denote by *j* the size of one class, 1 ≤ *j* ≤ *m −* 2, and then the size of the other class is *m −* 1 *− j*, 1 ≤ *m −* 1 *− j* ≤ *m −* 2. A triplet has *i*_1_ as one of the leaves and *i*_0_ as its center if and only if the two other leaves are in different classes. Thus, the number of such triplets is *j*(*m −* 1 *− j*), which is greater than or equal to *m −* 2. ▪

**Fig. 14.**
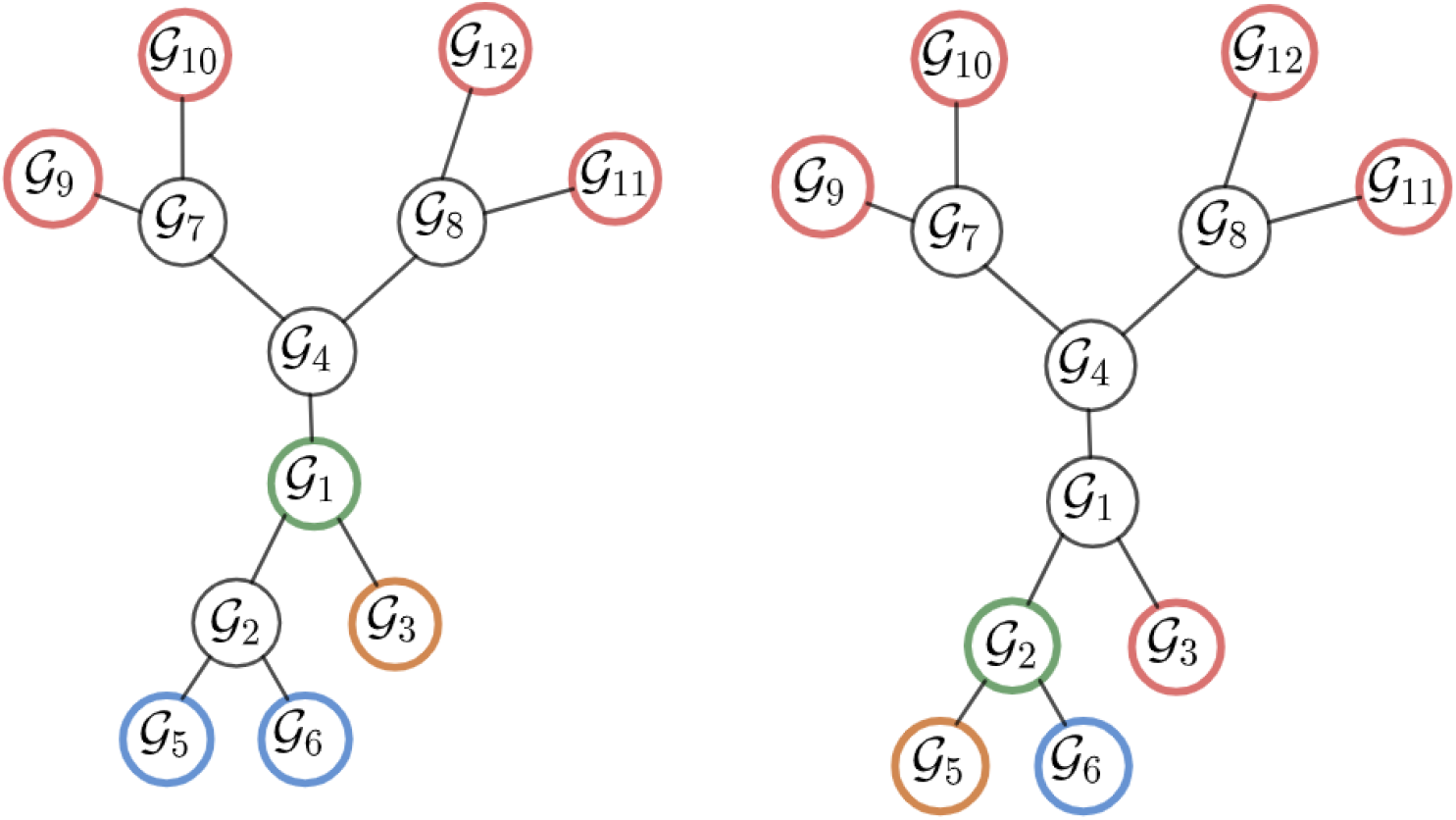
Estimating Directional Lengths of Edges Connected to Leaves. (L) To estimate directional length for the edge connected to 𝒢_3_, we may use any one of the triplets whose center is 𝒢_1_. Such triplets consist of a blue leaf, a red leaf, and 𝒢_3_. The number of such triplets is the number of blue leaves times the number of red leaves-a total of eight triplets. (R) 𝒢_5_ belongs to a cherry. In this case, we can use only five triplets to estimate directional lengths for the edge connected to 𝒢_5_. In this tree, for any leaf and for all the triplets it belongs, if we compute the directional length to the center of the triplet and compute the mean of the five smallest, we obtain an estimator for the directional length of the edge connected to the leaf.

**Fig. 15.**
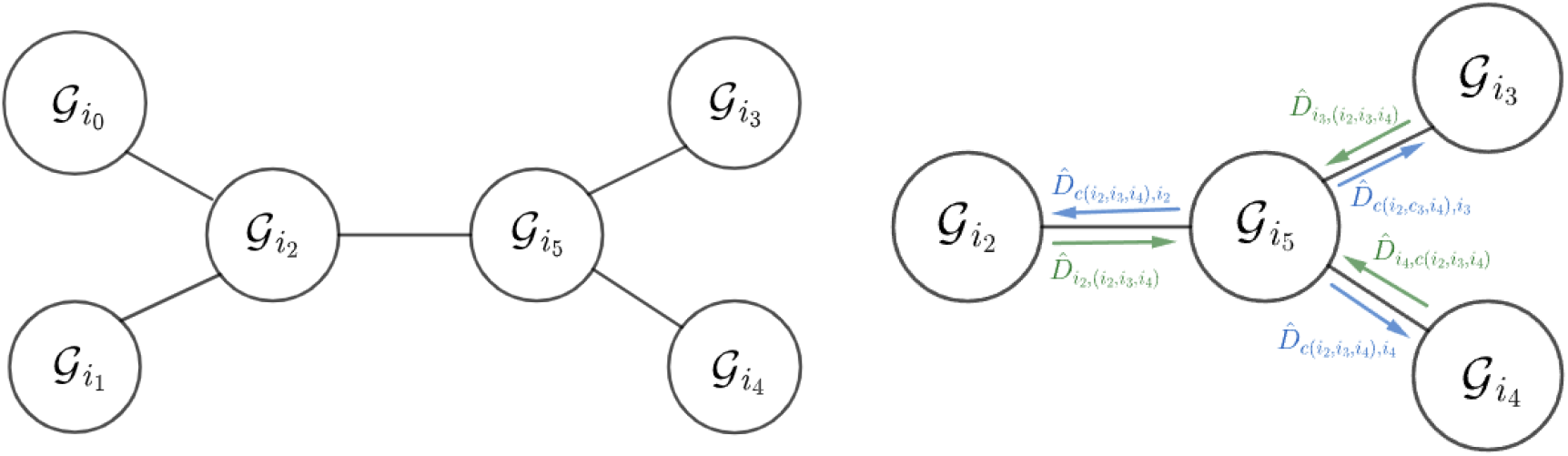
Quartet of leaves containing the chosen Cherry: (L) 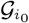 and 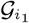 are the cherry sisters, 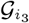 and 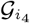 are any two other leaves. 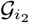 is the parent of the cherry. 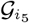 is the center of 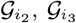 and 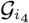.(R) For each such quartet of leaves, we estimate the directional distances of the triplet 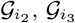 and 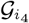.

We now provide a formal definition for the estimators 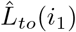 and 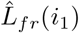 corresponding to leaf *i*_1_. Let 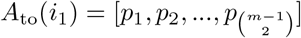, be a 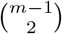 - long list corresponding to all pairs of distinct leaves without *i*_1_, such that 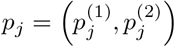 i.e. for any 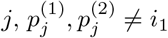, and assume 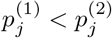.In addition, it is assumed that *A*_to_(*i*_1_) is sorted by the distance of the center 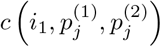 to *i*_1_ so that if *j*_1_ *< j*_2_ then the pairs 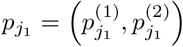 and 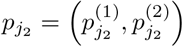 satisfy

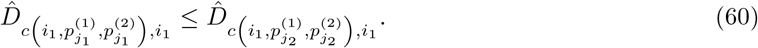

We now define an estimator for the directional distance *L*_*to*_(*i*_1_);

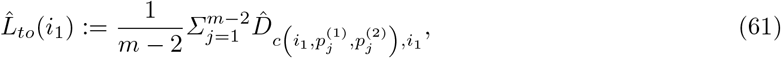

where the sum is over the first (i.e. minimal) elements in the sorted list 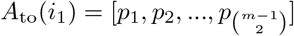. Similarly if the pairs 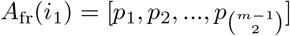 have an order that satisfies for *j*_1_ *< j*_2_

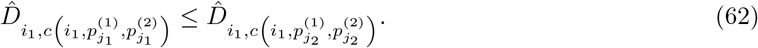

The estimator for *L*_*fr*_(*i*_1_) is,

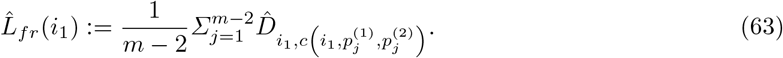

The next theorem deals with the consistency of 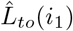 and 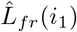.

#### Observation 10

*the estimators* 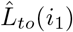 *and* 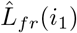 *are consistent such that if N*_0_ → ∞ *then*,

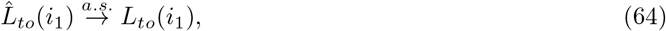

*and*

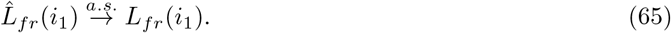

**Proof:** by definition *i*_0_ is the neighbour *i*_1_, thus it is the closest possible center to *i*_1_ (in both directions), hence, by observation 8 for any *j* such that 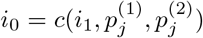 then

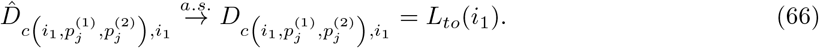

Otherwise, if *i*_0_ is not the center of the triplet 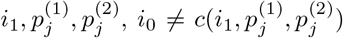 then the distance to the center is larger than the distance to *i*_0_ and,

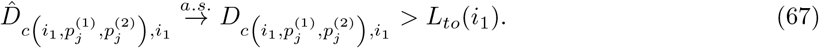

Therefore, if *N*_0_ → ∞ with probability 1 there is *K* such for any 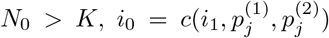 for 1 ≤ *j* ≤ *m −* 2. Thus, all the *m −* 2 smallest distances to *i*_1_ from centers are from *i*_0_ as a center. It follows that the components of the mean (Eq. 61) converge a.s. to *L*_*to*_(*i*_1_) and so does the mean. This completes the proof of the first part of the observation. The proof of the second part is similar. ▪

### B.3 Locating a Cherry

(Extended version of subsection 3.3)

The algorithm presented is of cherry-picking type, similar to the neighbor-joining algorithm [74] in that a pair of leaves *a, b*, supposed to be a cherry, is sought in *T*_*u*_. Then, these two leaf sisters are removed from *T*_*u*_. By doing so, in the new tree 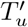, a subtree of *T*_*u*_, the sister’s parent node becomes a leaf, and the total number of leaves is reduced by one. We show how to repeat this procedure, locating cherries and removing their leaves, continuing to reduce the number of leaves, and, while doing so, assuming that the cherry-picking is correct, the original topology of *T*_*u*_ is reconstructed.

Here, we present our method for choosing such a cherry based on the estimators of directional distances for triplets of leaf genomes. Let *i*_1_ be a leaf, *i*_0_ be the (internal) node neighboring *i*_1_ and let *i*_2_ be another leaf. For the task of detecting a cherry pair, we define by *H*_*to*_ (*i*_1_, *{i*_1_, *i*_2_*}*) the mean of the distances from the centers of the triplets to *i*_1_ for all triplets *S* such that *i*_1_, *i*_2_ ∈ *S*. Similarly, *H*_*fr*_ (*i*_1_, *{i*_1_, *i*_2_*}*) is the mean distance from *Ĥ* to denote the means of the estimators of the correspondent distances.

We show later (Obs.12) that *H*_*to*_(*i*_1_, *{i*_1_, *i*_2_*}*) = *L*_*to*_(*i*_1_) if and only if the pair *{i*_1_, *i*_2_*}* forms a cherry, and similarly for *H*_*fr*_(*i*_1_, *{i*_1_, *i*_2_*}*) and *L*_*fr*_(*i*_1_). This is the key property for detecting a cherry.

Formally, if *i*_3_, …, *i*_*m*_ are the rest of the leaves, then we define

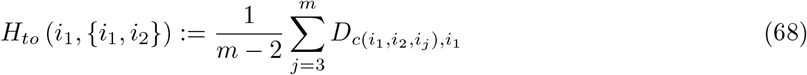

and similarly

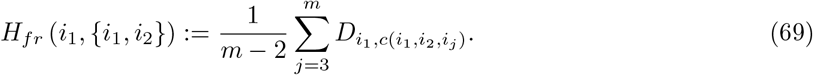

The following observation characterizes cherries;

#### Observation 11

*The pair i*_1_, *i*_2_ *forms a cherry if and only if i*_0_ *is the center of all triplets S such that i*_1_, *i*_2_ ∈ *S*.

As an immediate result of this observation, we have one more observation;

#### Observation 12

a. *The pair i*_1_, *i*_2_ *form a cherry if and only if H*_*to*_ (*i*_1_, *{i*_1_, *i*_2_*}*) = *L*_*to*_ (*i*_1_)
b. *If the pair i*_1_, *i*_2_ *is not a cherry, then H*_*to*_ (*i*_1_, *{i*_1_, *i*_2_*}*) *> L*_*to*_ (*i*_1_).
c. *The pair i*_1_, *i*_2_ *form a cherry if and only if H*_*fr*_ (*i*_1_, *{i*_1_, *i*_2_*}*) = *L*_*fr*_ (*i*_1_)
d. *The pair i*_1_, *i*_2_ *is not a cherry then H*_*fr*_ (*i*_1_, *{i*_1_, *i*_2_*}*) *> L*_*fr*_ (*i*_1_).

Naturally, we define estimators for *H*_*to*_ and *H*_*fr*_;

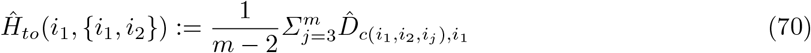

And

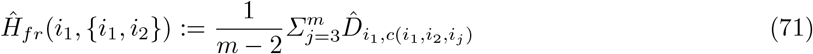

The following deals with the consistency of *Ĥ* ;

#### Observation 13

*Ĥto*(*i*_1_, *{i*_1_, *i*_2_*}*) *and Ĥf r*(*i*_1_, *{i*_1_, *i*_2_*}*) *are consistent estimators for H*_*to*_(*i*_1_, *{i*_1_, *i*_2_*}*) *and H*_*fr*_(*i*_1_, *{i*_1_, *i*_2_*}*) *respectively with almost surely convergence*.

**Proof:** The consistency and almost surely convergence of *Ĥto* and *Ĥf r* follows directly from the consistency and almost surely convergence of 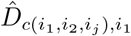 and 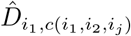.

We have one more observation on the relation between *Ĥ* and 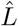;

#### Observation 14

a. 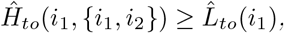.
b. 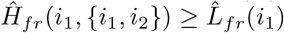.

**Proof:** The estimator 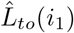 consists of the definition of the smallest distances *m−* 2 from the center to *i*_1_. The estimator 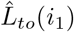 consists of *m−* 2 distances from the centers that are not necessarily the smallest. Hence, the result (a.) follows. The reasoning for (b.) is similar. ▪

The subtruction of the functions *Ĥ* and 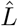 give positive functions that converge almost surely to zero only for cherries and to a positive limit for pairs that are not cherry;

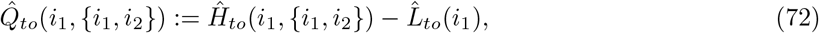

and

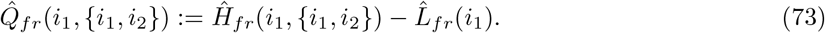

Then 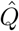 have the desired properties;

#### Observation 15

a. *By observation 14 above it follows that* 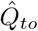 *and* 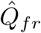 *are nonnegative*.
b. *Observations 10, 12 and 13 mean that if N*_0_ → ∞ *and the pair i*_1_,*i*_2_ *form a cherry, then* 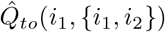 *and* 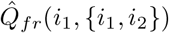 *almost surely converge to zero. Similarly, if N*_0_ → ∞ *but the pair i*_1_,*i*_2_ *is not a cherry, then* 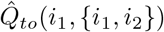 *and* 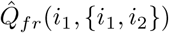 *almost surely converge to a positive number*.

Each pair of leaves *i*_1_ and *i*_2_ is graded using four *Q* values;

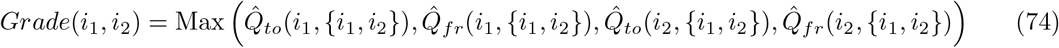

#### Observation 16

*From the almost surely convergence in the previous observation, it follows that in probability* 1 *there is K such that if N*_0_ *> K then the choice of minimized Grade is correct*.

The pair of leaves *a, b* that is chosen as cherry and in the reconstructed tree minimizes the function *Grade*. The directed distances 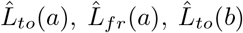 and 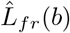 are the distances of the edges of the leaves in the reconstructed tree. Let *c* be the node in *T* connected to *a* and *b*. Then by Eq. 26 we estimate for *N*_*c*_ by geometric mean of two estimations:

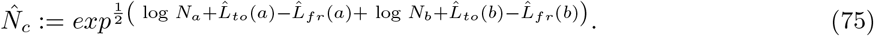

### B.4 Reducing the Number of Leaves by One

(Extended version of subsection 3.4)

Denote by *i*_0_, *i*_1_ the indexes of the cherry sisters. Let us denote by *i*_2_ their parent index. Let 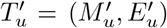 be the subtree of *T*_*u*_ after the sister leaves were removed. To continue with 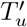 the same as we did with *T*_*u*_, we first need to estimate the directed distances of the triplets of leaves with *i*_2_ as one of the leaves. It was assumed that *T*_*u*_ has four or more leaves, so let *i*_3_ and *i*_4_ be any two other leaves of the tree *T*_*u*_. The following is how we estimate the directed distances of the triplet *i*_2_, *i*_3_, *i*_4_ from 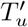 based on the triplets *{i*_0_, *i*_3_, *i*_4_*}, {i*_1_, *i*_3_, *i*_4_*}, {i*_0_, *i*_1_, *i*_3_*}, {i*_0_, *i*_1_, *i*_4_*}* and the directed distances of the leaf edges of *i*_0_ and *i*_1_ from *T*_*u*_. Using the estimators shown in Fig. 16. This is done as follows: The next four definitions are each a mean between two estimators for the same distance, under the assumption that *i*_0_ and *i*_1_ form a cherry,

**Fig. 16.**
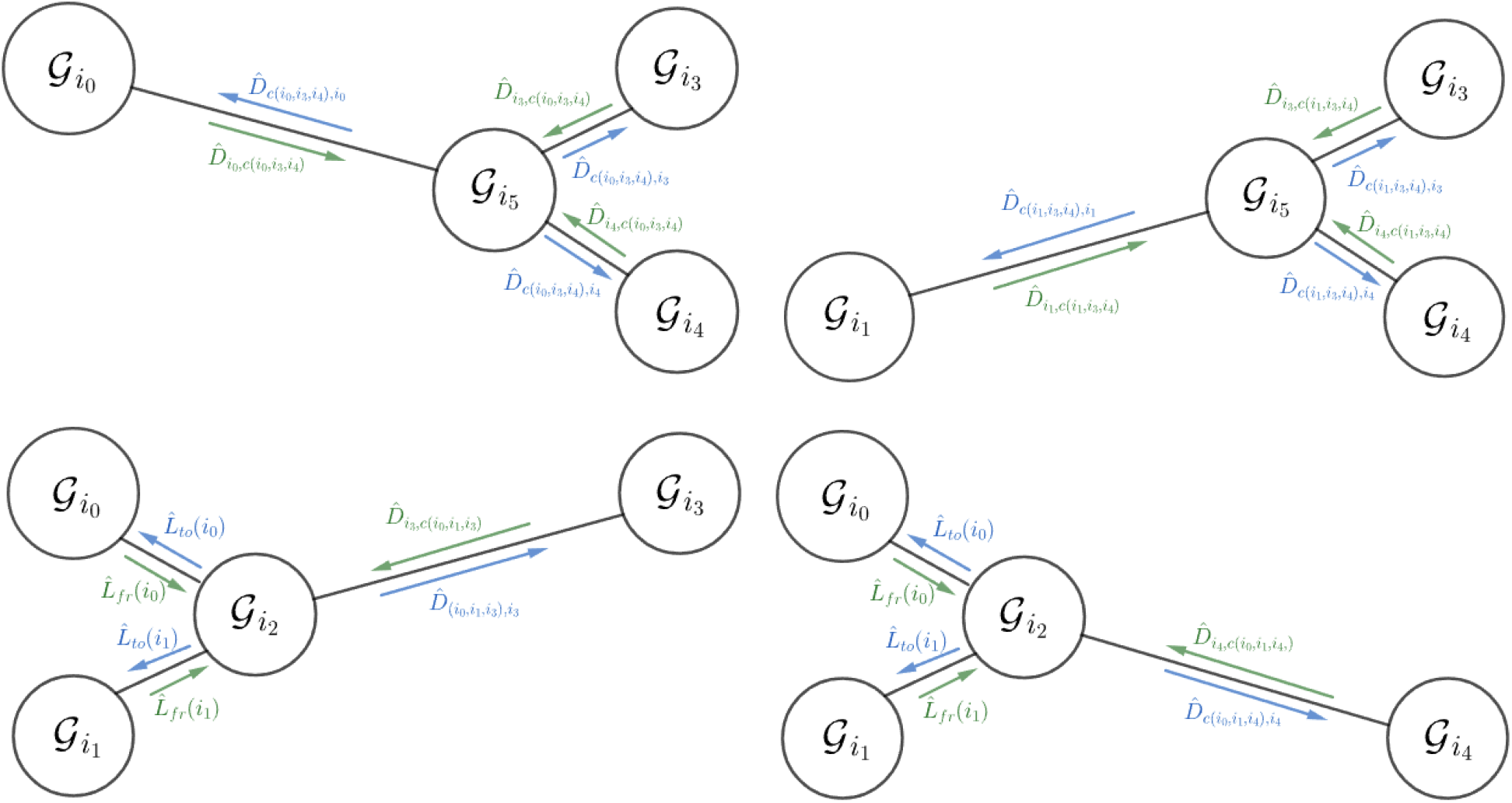
The estimators we use to estimate the directional distances of the triplet. 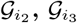 **and** 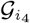.

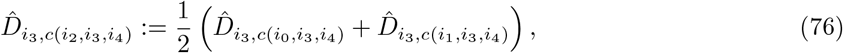

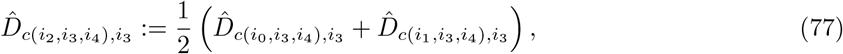

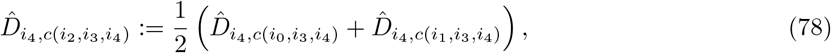

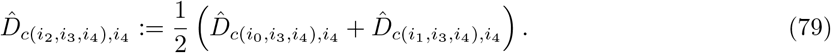

#### Corollary

From the consistency of the triplet directed distances and the previous observation we deduce the consistency of Eq. 76-79.

With the same assumption that *i*_0_ and *i*_1_ form a cherry, the following two definitions are each mean of four estimators for the same distance,

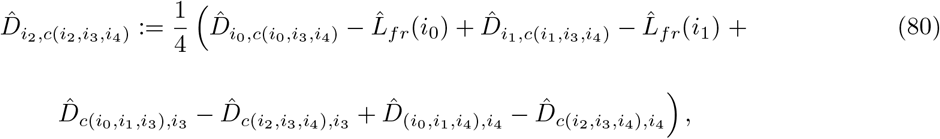

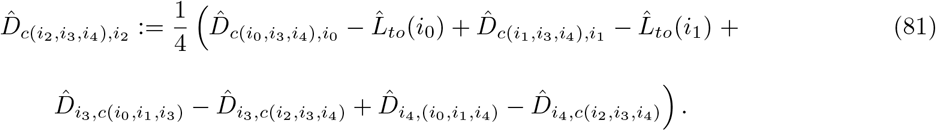

##### Observation 17

*The estimators in Eq. 80- 81 are consistent*.

**Proof:** we show the proof for Eq. 80. The proof for 81 is similar. If *i*_0_ and *i*_1_ are cherry, then 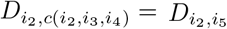. Because of the additivity,

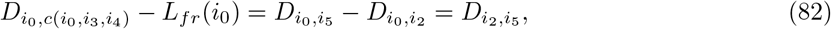

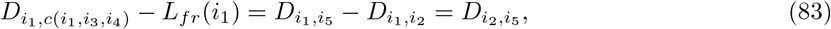

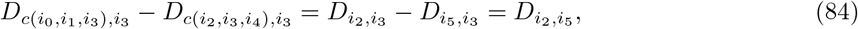

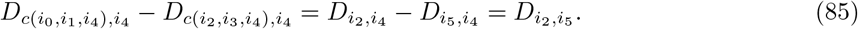

Thus, because of the consistency of the components of the mean that all tend to 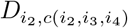 and because of the previous observation, the proof follows. ▪

These estimators are calculated where *i*_3_ and *i*_4_ run over all pairs of leaves that do not equal *i*_0_ or *i*_1_. This gives all the estimators that are needed to pick the next cherry in 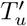.

### B.5 The End of the Algorithm

At the end of the loop there is only one triplet with three leaves and the directed distances from the center to the leaves of the triplet and vice versa. The center of the triplet is the last node that is added to the reconstructed tree, and if the triplet is *a, b, c* and the center is *d* then by Eq. 26 we estimate for *N*_*d*_ by geometric mean of three estimations:

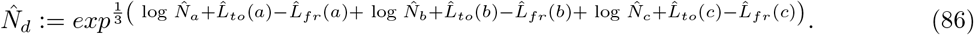

### B.6 The Formal TDDR Algorithm

The following is the pseudocode of the algorithm:

#### Algorithm 2

TDDR

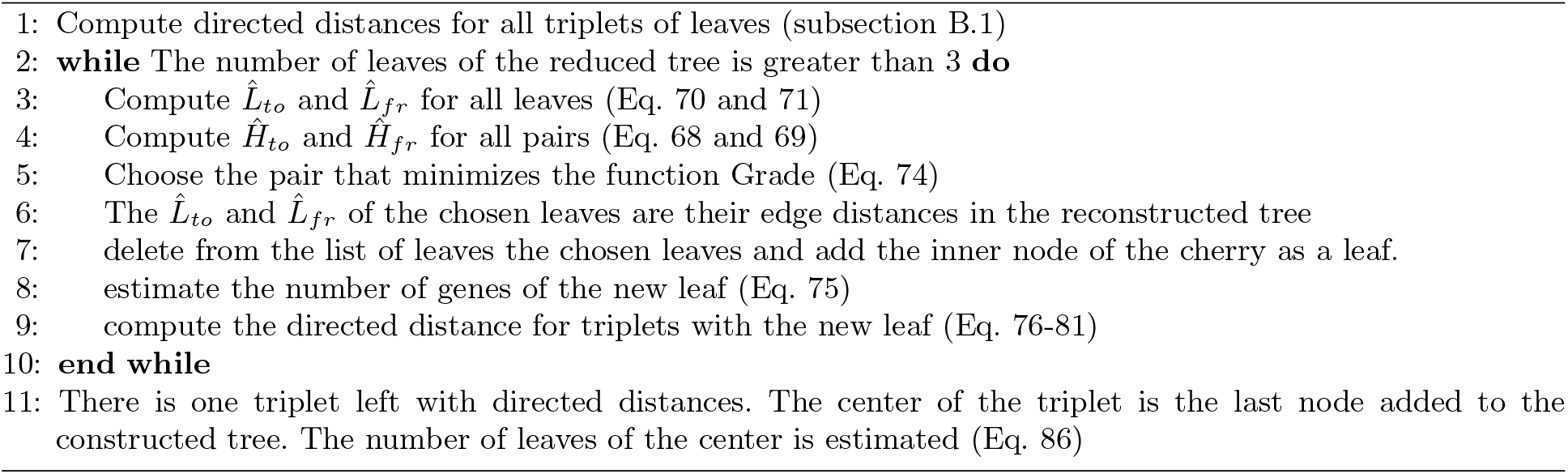

Assume that the data consist of a family of *m* leaf genomes and *n* is the maximum number of genes per genome, then the complexity of the first part (step 1) is *O*(*m*^3^*n*) due to the intersections that are computed for each triplet of genomes. The complexity of the second part (steps 2-11) is *O*(*m*^4^ log *m*) which is due to the computation of 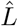 which consists of sorting a list of 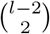 items for each of *l* leaves, where *l* goes from *m* to 3. Therefore, the complexity of the algorithm is *O*(*m*^3^*n* + *m*^4^ log *m*).

### B.7 Example of Tree Reconstruction Algorithm for a Tree with Four Leaves

In this subsection, we demonstrate a reconstruction of a tree with four leaves. The true tree with which we work appears in Fig. 17(L). The unrooted tree that we aim to reconstruct appears on the right. The edges of the rooted tree represent TRPs whose parameters (as directed distances) were taken randomly from the exponential distribution with an expectancy of 0.05. These parameters appear in Fig. 18(L). We produced genomes from the root down the tree (using the tree parameters) to obtain the leaf genomes. We continue the example according to the algorithm steps.

**Fig. 17.**
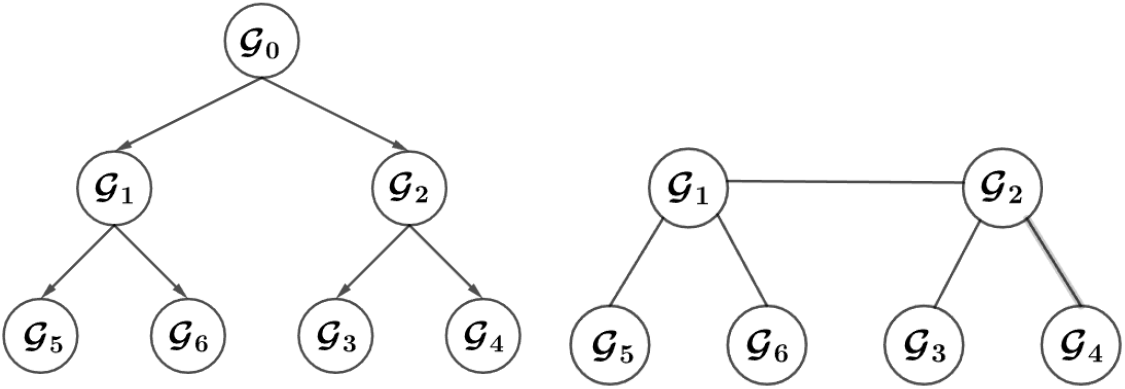
(L) Random tree with four leaves. (R) The unrooted tree that should be reconstructed.

**Fig. 18.**
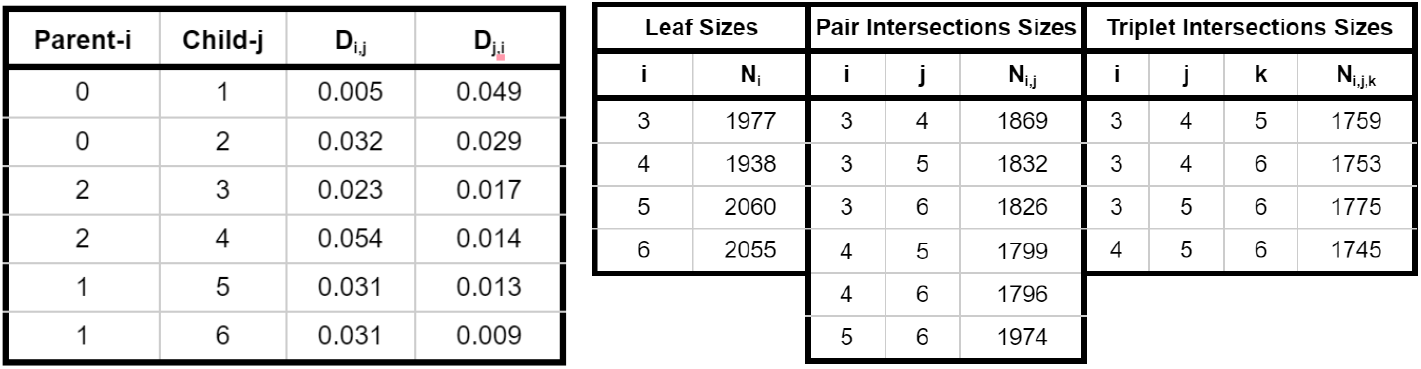
(L) Parameters: The directed distances of the tree edges are parameters to be estimated. **(R) Data: The data we use are the intersection of pairs and intersections of triplets of leaves**.

**step 1** The leaf genome sizes and their pairwise and tripletwise intersections are computed. For the example, the summary appears in Fig. 18(R). These data are all that is needed for tree reconstruction. For each triplet of leaves, we reconstruct an unrooted tree of three leaves, i.e., we compute the directed distances from the leaves to the center and vice versa. For example, according to equations 39-42, with the data from Fig. 18(R):

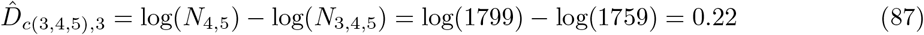

and

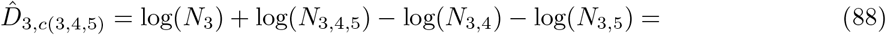

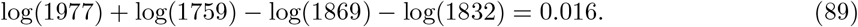

The directed distances of the example appear in Figure 19.

**Fig. 19.**
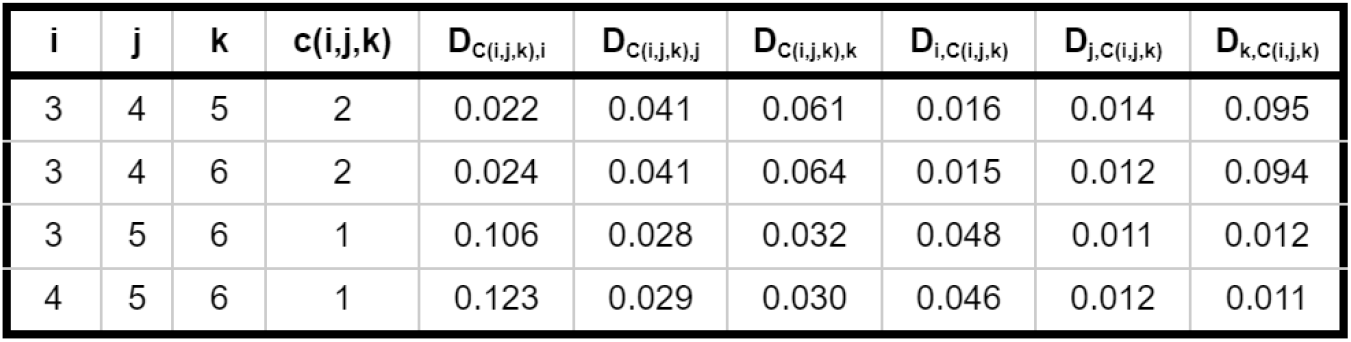
Estimates for triplets directed distances: At the beginning of the algorithm, the directed distances (from each leaf to the center of the triplet and vice versa) are calculated for all triplets of leaves.

**step 3:** For each leaf, the directed distances of its edge 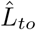 and 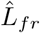 are estimated and presented in Fig. 20 (L). For example, to compute 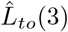 (see Eq 61) we compute the mean of the two (from three) smallest distances from centers to leaf 3 in Figure 19:

**Fig. 20.**
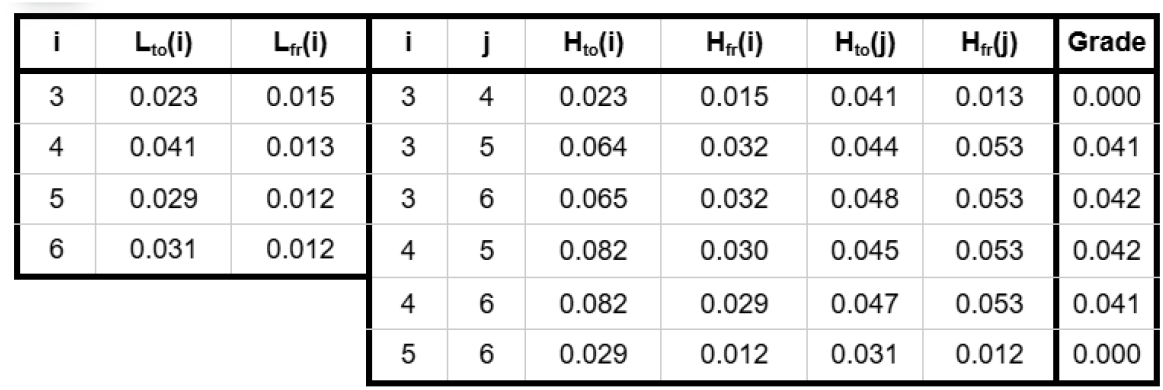
The columns *L*_*to*_(*i*) and *L*_*fr*_(*i*) are estimators for the edge directed distances of leaf *i*. If *i, j* form a cherry the values in columns *H*_*to*_(*i*), *H*_*fr*_(*i*), *H*_*to*_(*j*), *H*_*fr*_(*j*) tends to the correspondent L values. Otherwise, they are larger than the correspondent L values. The Grade column is calculated based on the H and L values. The pair that is chosen is the one that has the smallest Grade.

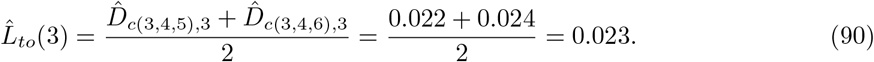

Similarly, 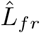 (3) (see Eq. 63) is the mean of the two (from three) smallest (from three) distances from leaf 3 to centers:

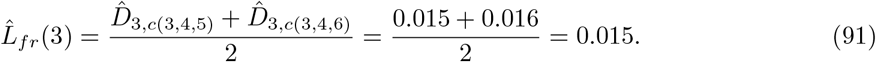

**step 4:** For each pair of leaves, *Ĥto* and *Ĥf r* are estimated and presented in Fig. 20 (M). By Eq. 61, for the pair 3, 4, *H*_*to*_(3, 3, 4) = *L*_*to*_(3) and *H*_*fr*_(3, 3, 4) = *L*_*fr*_(3). for the pair 3, 5

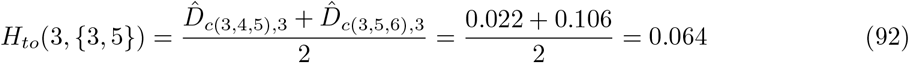

and by Eq 71,

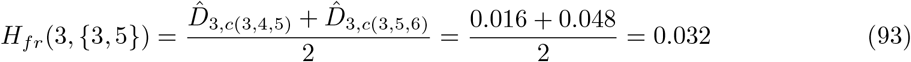

**step 5:** For each pair of leaves, We compute Grade and choose as the cherry the pair that minimizes Grade. The Grade for each pair is presented in Fig. 20 (R). For example, for the pair 3, 5 by equation 74:

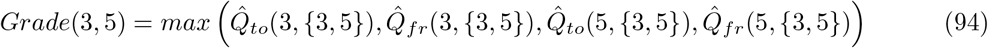

and by equations 72, 73, 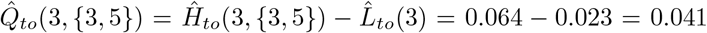, similarly,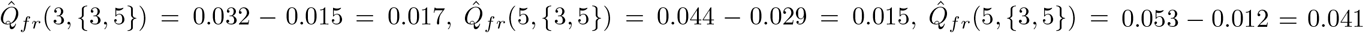. Hence *Grade*(3, 5) = 0.041. Two pairs 3, 4 and 5, 6 achieve the minimum of *Grade* = 0. Here, we pick 3, 4 as the cherry to continue with the example.

**step 6:** The directed distances 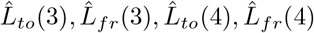 are the edge distances in the reconstructed tree.

**step 7:** The leaves of the cherry are omitted from the leaf list, and the inner node (2) of the cherry is now considered a leaf.

**step 8:** The size of *N*_2_ in the reconstructed tree is estimated

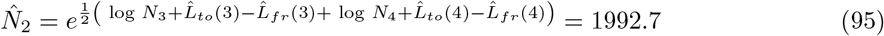

**step 9:** The directed distances for triplets that include the new leaf are computed. In this example, the pair 3, 4 is a real cherry pair and their real inner node is 2. In this example, we need to compute only the directed distances of the triplet 2, 5, 6 according to Eq. 76-81 substituting *i*_0_ = 3, *i*_1_ = 4, *i*_2_ = 2, *i*_3_ = 5, and *i*_4_ = 6.

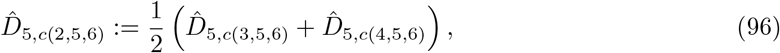

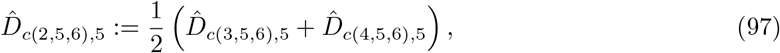

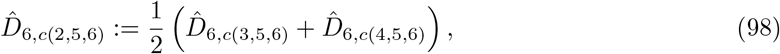

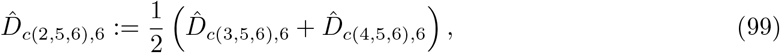

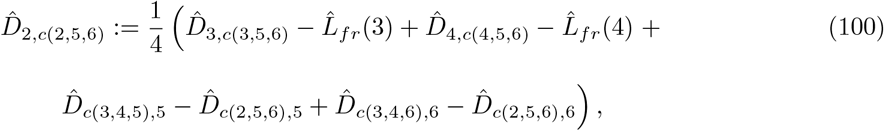

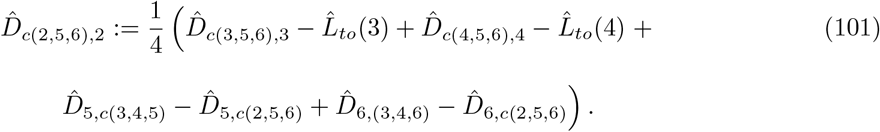

The results appear in Fig.21.

**Fig. 21.**
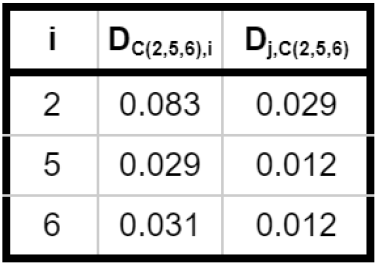
Updating Triplets distances for a new leaf: After choosing a cherry and replacing them by one leaf we calculate the directed distances for the new leaf triplets. In the case of this example, we need to calculate the distances for one triplet.

**step 11:** In this stage, at the end of the “while” loop, we are left with a tree with three leaves. The center of the triplet 2, 5, 6 is the node 1. estimation of *N*_1_

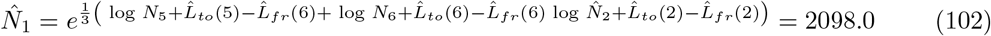

With the leaves 3, 4 that are connected to leaf 2, the unrooted tree in 17 is fully reconstructed with all edge directed distances (see Fig 22).

**Fig. 22.**
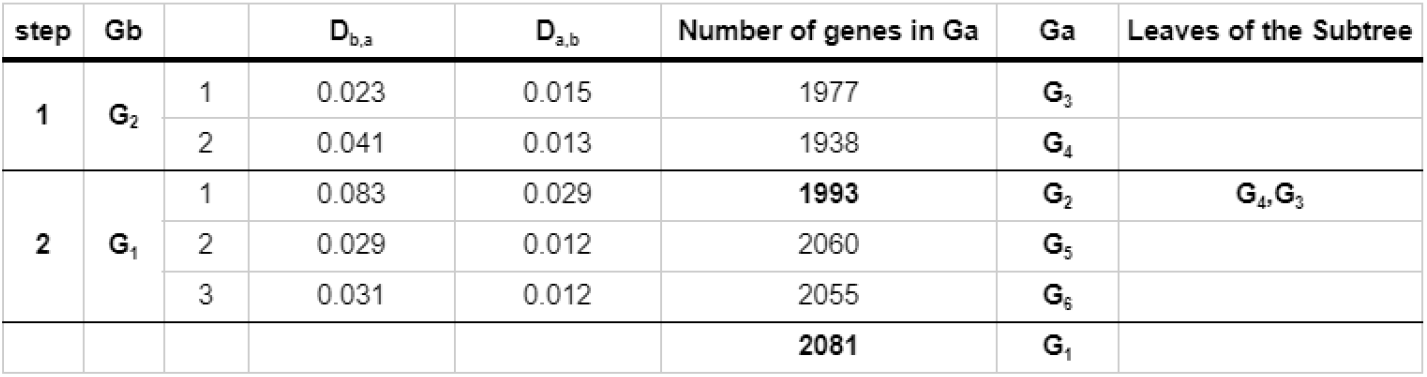
The example constructed tree: This is a description of the process of cherry pair picking and it is also the structure and estimates for the reconstructed tree. The column *G*_*a*_ is the column of the pairs of leaves that were picked. The column *G*_*b*_ is the new leaf that replaces the cherry sisters and it is an internal node in the reconstructed tree. The columns *D*_*b,a*_ and *D*_*a,b*_ are the edge-directed distances. The column *N*_*a*_ is the number of genes in *G*_*a*_. For the leaves *G*_3_ *− G*_6_, *N*_*a*_ is exact, and for the internal nodes *G*_1_ *− G*_2_ this is the estimated number of genes.

## C Experimental Results

(Extended version of Sect. 4)

### C.1 Simulations

#### Single Edge TRP Simulation

Here, we describe the TRP process that is simulated on a single edge. The random distances from the simulated TRP, *D*_0,1_ and *D*_1,0_, are chosen independently of an exponential distribution. We set the total time *T* = *D*_0,1_ + *D*_1,0_ (the length of the edge). The birth rate was set as 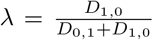 and the death rate as 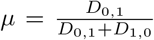. The parent is indicated by 𝒢 (0) and the child by 𝒢(*T*). The child is produced from the parent through a sequence of events. The genome created by the ith event is denoted by 𝒢 (*t*_*i*_). Each genome 𝒢 (*t*_*i*_) is represented by a sequence of *n*_*i*_ distinct genes represented by serial numbers. The genome 𝒢 (*t*_*i*+1_) is produced from 𝒢 (*t*_*i*_) as follows; a random time δ_*i*_ is chosen from exponential distribution with rate *n*_*i*_. if *t*_*i*_ + δ_*i*_ *> T* then the process is done and 𝒢 (*T*) = 𝒢 (*t*_*i*_). Otherwise, *t*_*i*+1_ = *t*_*i*_ + δ_*i*_, and in probability *λ*, the event is a birth event; otherwise, with probability *µ*, it is a death event. A random gene is eliminated from the sequence if it is a death event. If it is a birth event, a random gene is chosen and a new gene is added, with probability half next to the selected gene or probability half as the gene that precedes the chosen gene. To comply with the characteristics of the TRP, the new gene is belongs to the same *M*_*q*_ as the selected gene. The new gene has a new serial number that has not been used until now, even in other TRPs.

#### Simulations of Trees with Three Leaves

For simulations of trees with three leaves we start with 𝒢_0_ with a size of 2000 genes, then produce from the parent 𝒢_0_ by two independent TRPs the children’s genomes 𝒢_1_ and 𝒢_2_ and then from 𝒢_1_ as parents we produce again by two independent TRPs the children 𝒢_3_ and 𝒢_4_. We perform simulations for mean directional edge lengths 0.005, 0.01, 0.015, …, 1; 10, 000 repetitions for each length. The 0.05 and 0.95 quantiles of the produced genomes, *N*_1_, *N*_2_, *N*_3_ and *N*_4_ are presented in Fig. 23a. We estimate the directional distances for the estimators presented in Eqs. 39-44. The standard errors for estimators of the directional distances are shown in Fig. 23b. The errors for the different edges are close enough to justify the presentation of the combined error. We also estimate the size of the internal node 𝒢_1_ according to Eq. 54 and the standard errors are shown in Fig. 23c..

**Fig. 23.**
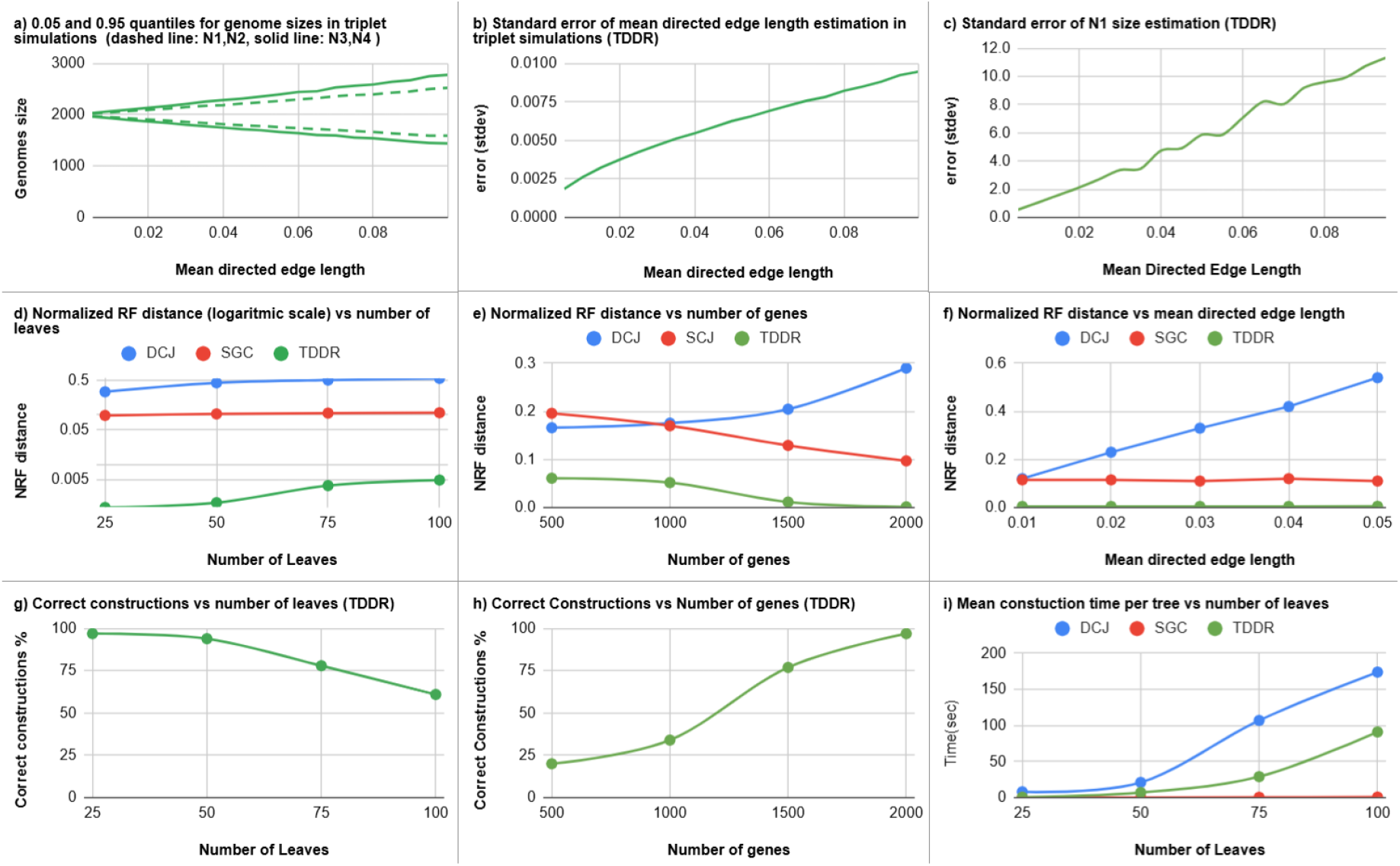
a) Sizes of genomes in simulations of trees with three leaves: for 10,000 repetitions and mean directed edge lengths 0.005 *−* 0.1, these are the 0.5 and 0.95 quantiles for *N*_1_ and *N*_2_ and for *N*_3_ and *N*_4_. Genome sizes *N*_1_ and *N*_2_ have the same distributions, as do *N*_3_ and *N*_4_. **b) Estimation error for the directed distances in simulations of trees with three leaves:** for 10,000 repetitions and mean directed edge lengths 0.005 *−* 0.1, standard errors for the directed distances *D*_1,2_, *D*_2,1_, *D*_1,3_, *D*_3,1_, *D*_1,4_ and *D*_4,1_. **c) The estimation standard error for** *N*_1_ **in simulations of trees with three leaves:** the size of 𝒢_1_ for 10,000 repetitions and mean directed edge lengths 0.005 *−* 0.1. **d) Normalized RF distance (logarithmic scale) vs number of leaves**. Mean directed edge length 0.05, Tree sizes: 25, 50, 75, and 100. For each size, 100 repetitions. **e) Normalized RF distance vs number of genes (size of** *N*_0_ ). Mean directed edge length 0.05, Tree size is 25 leaves, sizes of 𝒢_0_ : 500, 1000, 1500, and 2000. For each size, 100 repetitions. **f) Normalized RF distance vs mean edge length:** Tree size 100 leaves, mean directed edge lengths 0.01-0.05. For each mean length, 100 repetitions. **g) Percentage of correct reconstructed trees vs number of leaves**. Mean directed edge length 0.05, Tree sizes: 25, 50, 75, and 100. For each size, 100 repetitions. **h) Percentage of correct reconstructed trees vs number of genes**. Mean directed edge length 0.05, Tree size is 25 leaves. For each size, 100 repetitions. **i) Mean CPU time for one repetition vs number of leaves**. Mean directed edge length 0.05. Tree sizes: 25, 50, 75, and 100. For each size, 100 repetitions.

#### Simulations for the Tree Reconstruction Algorithm

The algorithm (TDDR) is described in Sect. B. In each simulation repetition, a new (random) rooted tree is drawn starting with the genome 𝒢_0_, which has *N*_0_ genes at the root. Genome evolution proceeds recursively, starting from the root genome: A random leaf is chosen from the tree and two new child leaves are connected. Two independent TRPs create the genomes of these children of the new parent. This process continues until the desired number of leaves is reached. The leaf genomes resulting from this process are passed as input to the various packages under test. The TDDR is compared here with two different methods to create distance matrices implemented in the following packages;

Seq. 1 The trees have sizes of 25, 50, 75, and 100 leaves with a mean directed edge length of 0.05. For each tree size, we do 100 repetitions.

Seq. 2 Mean directed edge lengths 0.1, 0.2, 0.3, 0.4, 0.5 with trees of size 100 leaves. For each mean length of the edge, 100 repetitions.

Seq. 3 First genome *G*_0_ with sizes 500, 100, 1500, 2000 with trees of size 25 leaves. For each size, 100 repetitions.

For all methods, the reconstructed trees are tested against the true trees using the normalized Robinson-Foulds distance implementation from the ETE3 package (Fig. 23d-f). We also tested for TDDR the correct reconstruction rates for Seq.1 and Seq.3 (Fig. 23g-h) and mean time for tree reconstruction for the three methods (Fig. 23i).

### C.2 Real Data

(Extended version of Sect. 4.2)

The Alignable Tight Genomic Clusters (ATGCs) database [39] contains several million proteins from thousands of genomes organized into hundreds of clusters of highly closely related bacterial and archaeal genomes. Each ATGC is a taxonomy-independent cluster of two or more completely sequenced genomes. The ATGC database also includes many ATGC-COGs (Clusters of Orthologous Genes) and phylogenetic trees of the organisms within each ATGC. For the narrow phylogenetic spectrum of each ATGC, the probability of a duplication event inside an ATGC is very small [40,73]. In addition, orthology detection is significantly simplified compared to more general DB (e.g. EggNOG [29]). These facts make our model assumptions of infinite pangenome and a single copy COG highly realistic. We also note that no discordance between a gene-tree and the species-tree is possible under such a model. For the real data test, we used the following ATGCs; ATGC005, ATGC007, ATGC008, ATGC009, and ATGC032. The genome here is represented in SGC and TDDR as a set of distinct COGs (instead of genes). We used the same methods used for the simulations for tree reconstructions. We also used the phylogenetic tree downloaded from the ATGCs site. The various trees are compared by normalized RF and are shown in Figure 24. It can be seen that the differences between the directed distances on the same edge and between edges reflect real differences and support the need for the TRP model, untying gain and loss rates.

**Fig. 24.**
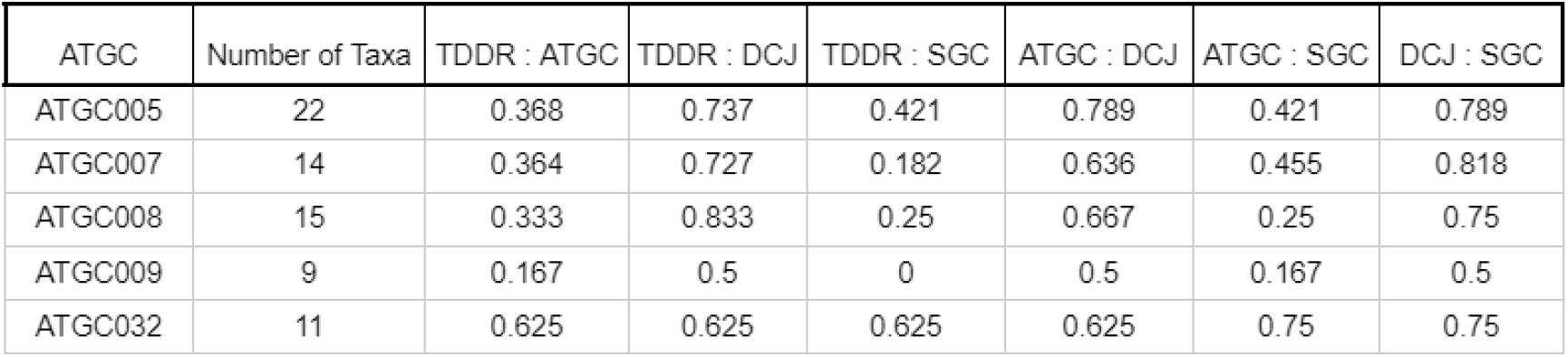
Normalized RF comparisons: The trees created by the three methods TDDR, DJC, SJC and the tree downloaded from the ATGCs site are compared by normalized RF distance, for the ATGCs: ATGC005, ATGC007, ATGC008, ATGC009 and ATGC032.

Fig. 25 shows the steps of the TDDR in constructing the ATGC009 tree. For each step in the column *G*_*a*_, one can find the two nodes for which the TDDR decides that they form a cherry and *G*_*b*_ is the node connected to the cherry. The direction distances *D*_*b,a*_ and *D*_*a,b*_ are given in two different columns. Considering the standard error of the algorithm demonstrated in simulations in Fig. 23b suggests that the differences between the directed distances on the same edge and between edges reflect real differences and support the need for our model.

**Fig. 25.**
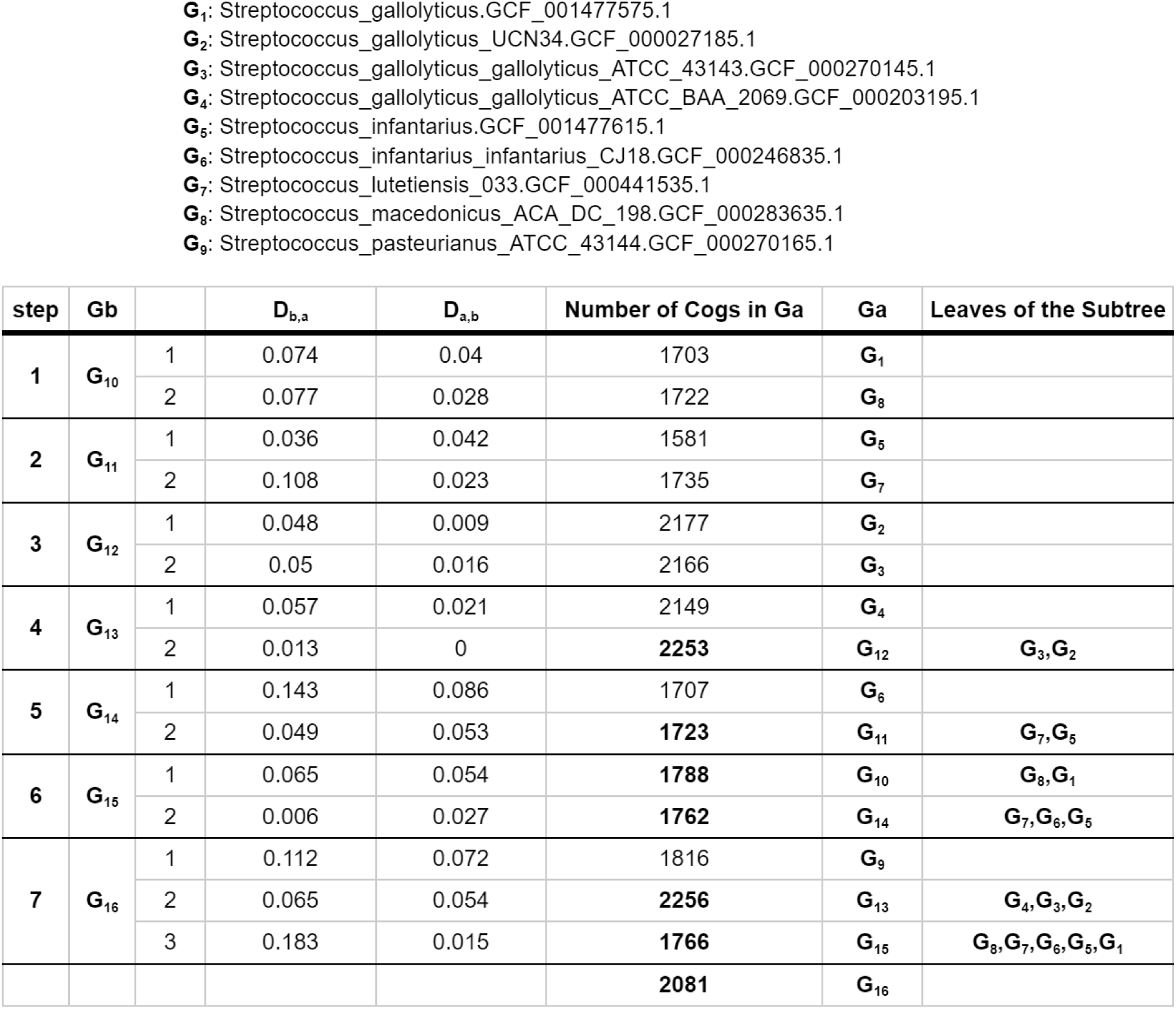
ATGC009 reconstruction: This is a description of the process of cherry pair picking and it is also the structure and estimates for the reconstructed tree. The column *G*_*a*_ is the column of the pairs of leaves that were picked. The column *G*_*b*_ is the new leaf that replaces the cherry sisters and it is an internal node in the reconstructed tree. The columns *D*_*b,a*_ and *D*_*a,b*_ are the edge-directed distances. The column *N*_*a*_ is the number of COGs in *G*_*a*_. For the leaves *G*_1_ *− G*_9_, *N*_*a*_ is exact, and for the internal nodes *G*_10_ *− G*_16_ this is the estimated number of COGs.

In each step, the table gives the number of cogs of the genomes in 𝒢_*a*_, and if the number is written in bold, it means it is estimated (see Eq. 75). The nodes of the subtree induced by the leaves 𝒢_1_, 𝒢_5_, 𝒢_6_, 𝒢_7_, 𝒢_8_ have less than 1800 cogs each, and the nodes of the subtree induced by the leaves 𝒢_2_, 𝒢_3_, 𝒢_4_ have more than 2150 cogs each. 𝒢_16_ and 𝒢_9_ are between 2081 and 1816, respectively.

The trees of ATGC009 are presented in figure 26. The TDDR and SGC have the same topology. They differ from the ATGC tree at the places of 𝒢_9_ which is connected to the TDDR tree at 𝒢_16_, and the cherry that is a cherry in all trees, 𝒢_1_ and 𝒢_8_ and its parent 𝒢_10_ and is connected to the TDDR tree 𝒢_15_ there connections nodes to the tree are replaced.

**Fig. 26.**
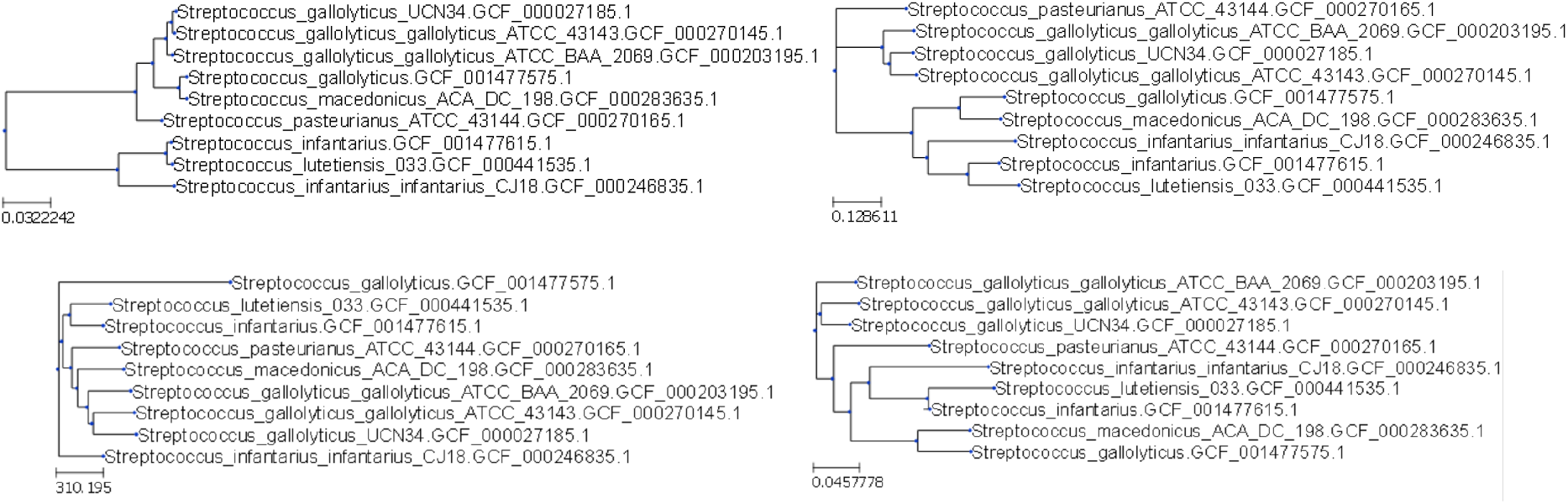
ATGC009. The upper left tree is the ATGC tree, the upper right tree is TDDR tree, the lower left tree is the DJC tree and the lower right tree is the SGC tree.

**Fig. 27.**
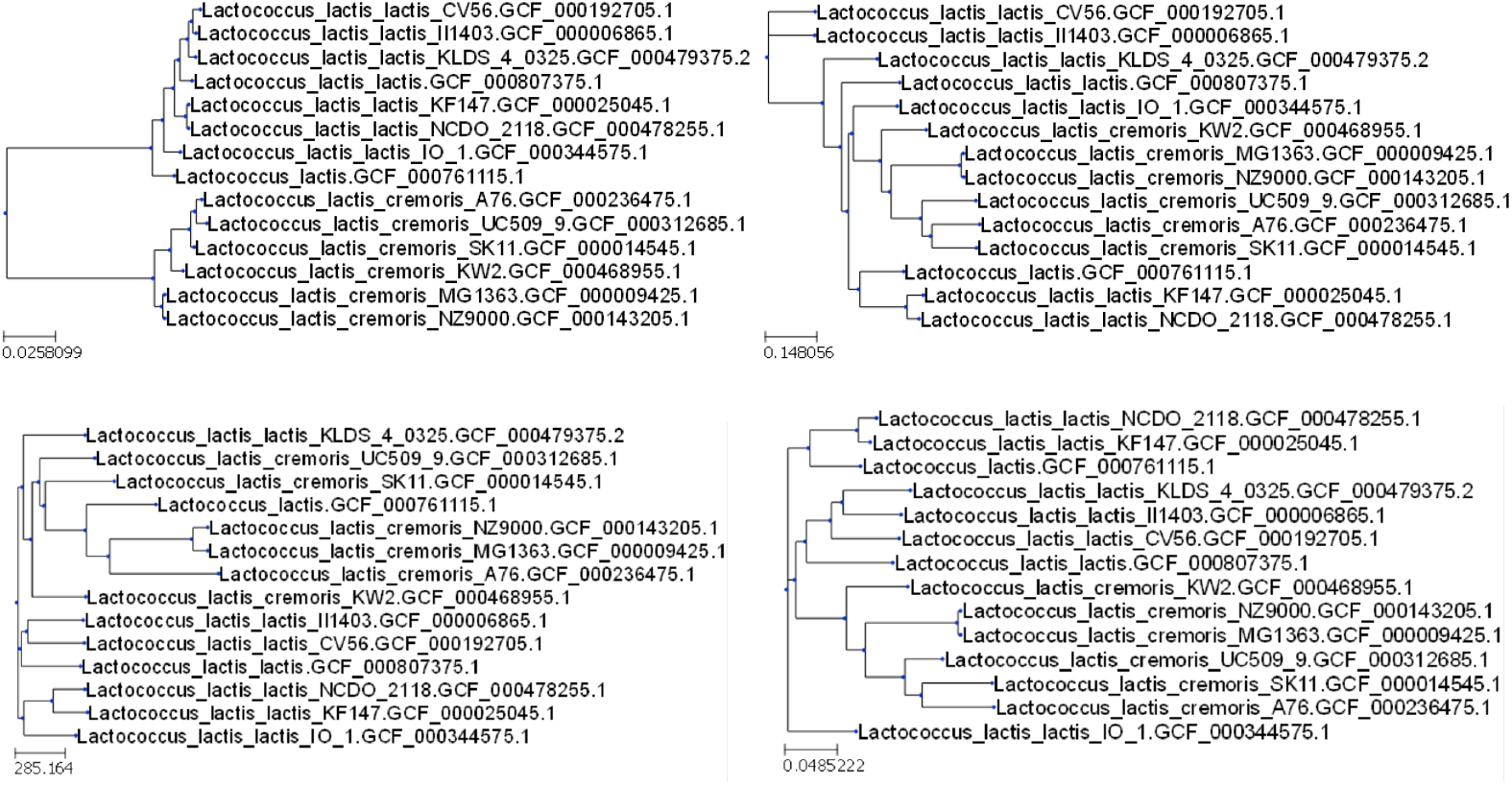
ATGC007. The upper left is the ATGC tree. The upper right is TDDR tree. Normalized RF distance with the ATGC tree: 0.36. The lower left is the DJC tree. Normalized RF distance with the ATGC tree: 0.64. The lower right is the SGC tree. Normalized RF distance with the ATGC tree: 0.45.

**Fig. 28.**
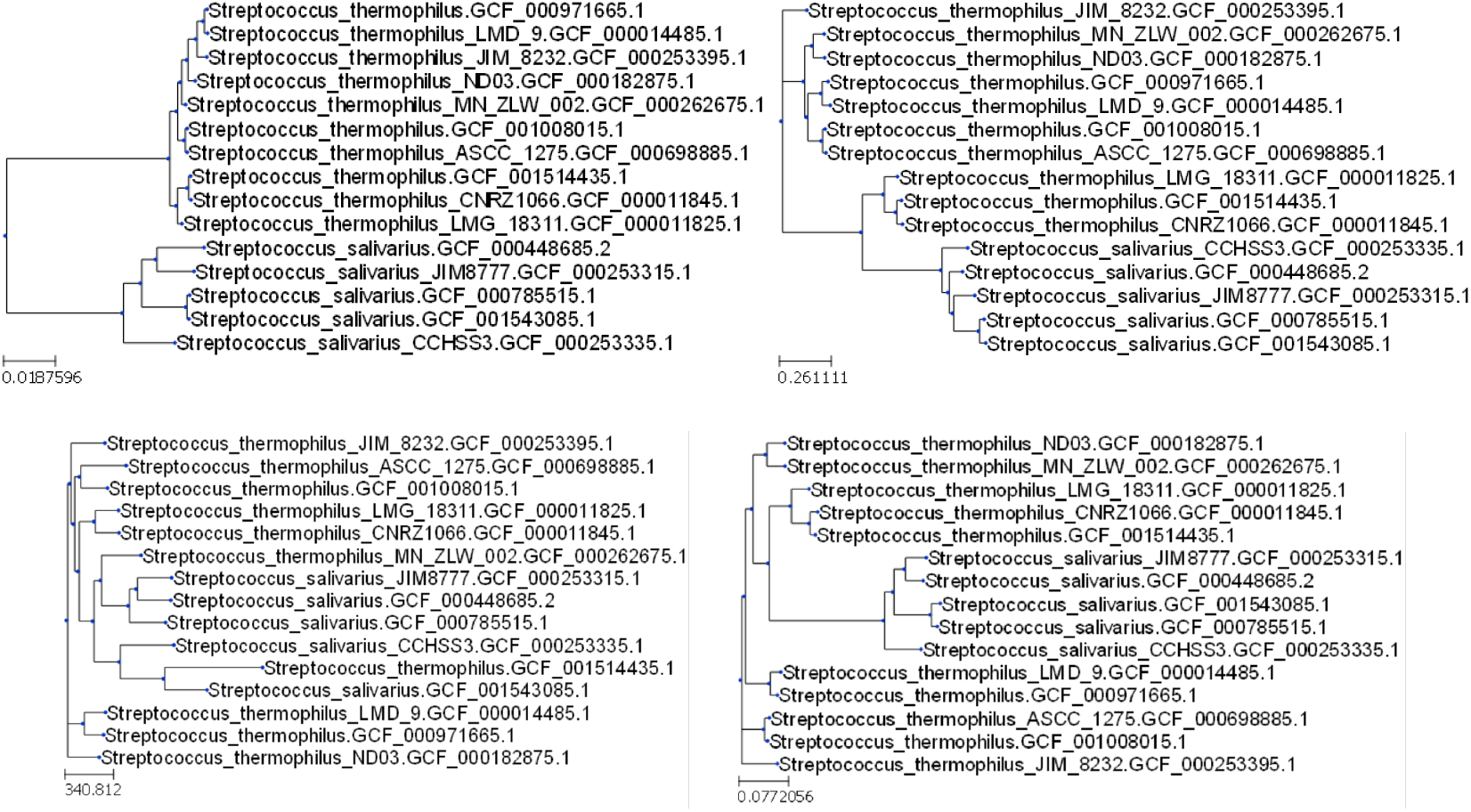
ATGC008. The upper left is the ATGC tree. The upper right is TDDR tree. Normalized RF distance with the ATGC tree: 0.33. The lower left is the DJC tree. Normalized RF distance with the ATGC tree: 0.66. The lower right is the SGC tree. Normalized RF distance with the ATGC tree: 0.25.

